# Heterochrony of axis segmentation underlies extreme morphogenesis in the Japanese eel

**DOI:** 10.64898/2026.04.29.721344

**Authors:** Ali Seleit, Kazuharu Nomura, Leon Hilgers, Michael Hiller, Kiyoshi Naruse, Yukinori Kazeto, Alexander Aulehla

## Abstract

Heterochrony is a major mode of vertebrate body plan evolution, yet its molecular and cellular basis remains poorly characterized. Here, we show that axis segmentation in *Anguilla japonica*—a species that forms 120 vertebrae—is driven by the temporal extension of somitogenesis. This is achieved through the prolonged maintenance of axial progenitors in the tail beyond the hatching stage, coupled to an extreme segment scaling regime that operates under the constraint of minimal axis growth. We identify delayed *Hox13* activation and sustained *Oct4* expression as molecular signatures of the prolonged segmentation program. Furthermore, we describe two spatially distinct axial progenitor pools that expand stemness in the tail. These findings reveal how the modulation of stemness in *time* and *space* drives extreme morphological evolution in vertebrates.

**One-Sentence Summary:** Spatiotemporal modulation of stemness drives extreme axis segmentation in the Japanese eel.

## Main Text

A hundred-fold difference in vertebral counts exists across vertebrates, ranging from just 7 in the frog *Paedophryne amauensis* (*1*) to over 750 in the slender snipe eel *Nemichthys scolopaceus* (*2*), making vertebral number one of the most striking features of body plan variation. The periodic addition of somites—the precursors of vertebrae—is highly conserved across vertebrates and is linked to the segmentation clock dynamics in the presomitic mesoderm (PSM) (*3, 4*). The modulation of the clock period, the duration of somitogenesis, or both, is thought to underlie species-specific segment numbers (*5, 6*). Yet the molecular, cellular and morphometric basis of this variation remains poorly characterized due to the limited tractability of vertebrate embryos outside a few well-established model organisms (*3, 7–9*).

To elucidate principles underlying vertebrate body plan variation, we leverage recent advances in aquaculture (*10*, *11*) to uncover the molecular and developmental mechanism underlying an extreme case of body axis segmentation in the Japanese eel, *Anguilla japonica,* a species with a highly complex life-cycle that forms up to 120 vertebrae (*12*, *13*). We show that the heterochrony underlying the expansion of body segments in *Anguilla japonica* is not driven by changes in the segmentation clock period but rather by a spatiotemporal extension of axial progenitors in the tail. This prolongation is coupled to an extreme segment scaling regime that operates under the constraint of minimal axis growth. We link this expanded segmentation program to key molecular and cellular mechanisms that extend stemness in *time* and *space*. First, we show that delayed *Hox13* cluster activation and sustained *Oct4* expression in the PSM maintain the stemness of the neuromesodermal progenitors (NMPs) beyond the hatching stage. Second, we discover two pools of NMPs exhibiting distinct temporal dynamics that spatially expand stemness in the tail. Our results highlight how spatiotemporal changes in stemness enable the modification of the axis segmentation program in vertebrates. Importantly, we reveal that the heterochronic changes in axis segmentation are uncoupled from overall developmental progression in the Japanese eel, demonstrating the modular control of development as the basis for body plan innovation.

### Extreme tissue scaling enables expanded segmentation under a minimal growth regime

To determine the morphometric basis of axial segmentation in *A. japonica*, we performed quantitative 3D volumetric analysis of nuclear-stained samples across somitogenesis stages (Fig. 1, A to E). We found a continuous reduction in PSM volume accompanied by a progressive decrease in somite size (Fig. 1, F and G; fig. S1, A to C; movie S1). By the 47-somite stage, newly formed somites are composed of only two cell layers along the anterioposterior axis, and subsequent somites undergo further reduction in size through compaction (fig. S1A). Unexpectedly, although the Japanese eel generates up to 120 somites, posterior axis growth is modest (3.1-fold) over the entire process (Fig. 1H; fig. S1, D to E; movie S2). This trajectory is reminiscent of zebrafish (*7*, *14*) but fundamentally distinct from the significant growth observed in mice and snakes (*5*, *8*, *14*). Analysis of the mitotic index (MI) in the unsegmented tail reveals an initial increase followed by a sharp decline (fig. S1, F and G), at no point does proliferation compensate for the loss of PSM tissue to somite incorporation. We show that the segmentation mode in *A. japonica* is characterized by a scaling regime in which the ratio of nascent somite size to PSM length is invariant over a large range (*i.e.* a 5-fold change in PSM length) (Fig. 1I). Importantly, the stable fraction of the PSM that gets incorporated into a segment is minimal, *i.e.* only 6.20% per cycle. As a result, despite having a comparable overall axis growth and PSM length as a zebrafish embryo (*7*, *14*), the eel PSM is segmented into 120 somites compared with 32 in zebrafish. This extreme form of scaling manifests most clearly when somites reach a minimal A-P length of two compacted cell layers (∼7 µm), each forming the anterior and posterior segment compartment, respectively (Fig. 1I; fig. S1 A and C). This demonstrates that the developmental constraint (*15*, *16*) on somite size is ultimately defined by a physical limit. To assess the ratio of abdominal to caudal somites in *A. japonica* we used the position of the anus and the anterior expression boundary of *HoxD12* as landmarks for the transition to caudal somite formation (*6*, *17*). Our data show that despite having ∼90 additional somites compared to medaka and zebrafish, the ratio of abdominal to caudal somites is conserved in all three species (fig. S1, H to K; fig. S2, A to D). This indicates that the extreme axis segmentation in eels results from a proportional expansion of both abdominal and caudal somite numbers (*17*, *18*). Collectively, our data reveal how a segmentation program with extreme tissue scaling permits segment number expansion under the constraint of minimal axis growth.

**Fig. 1.**
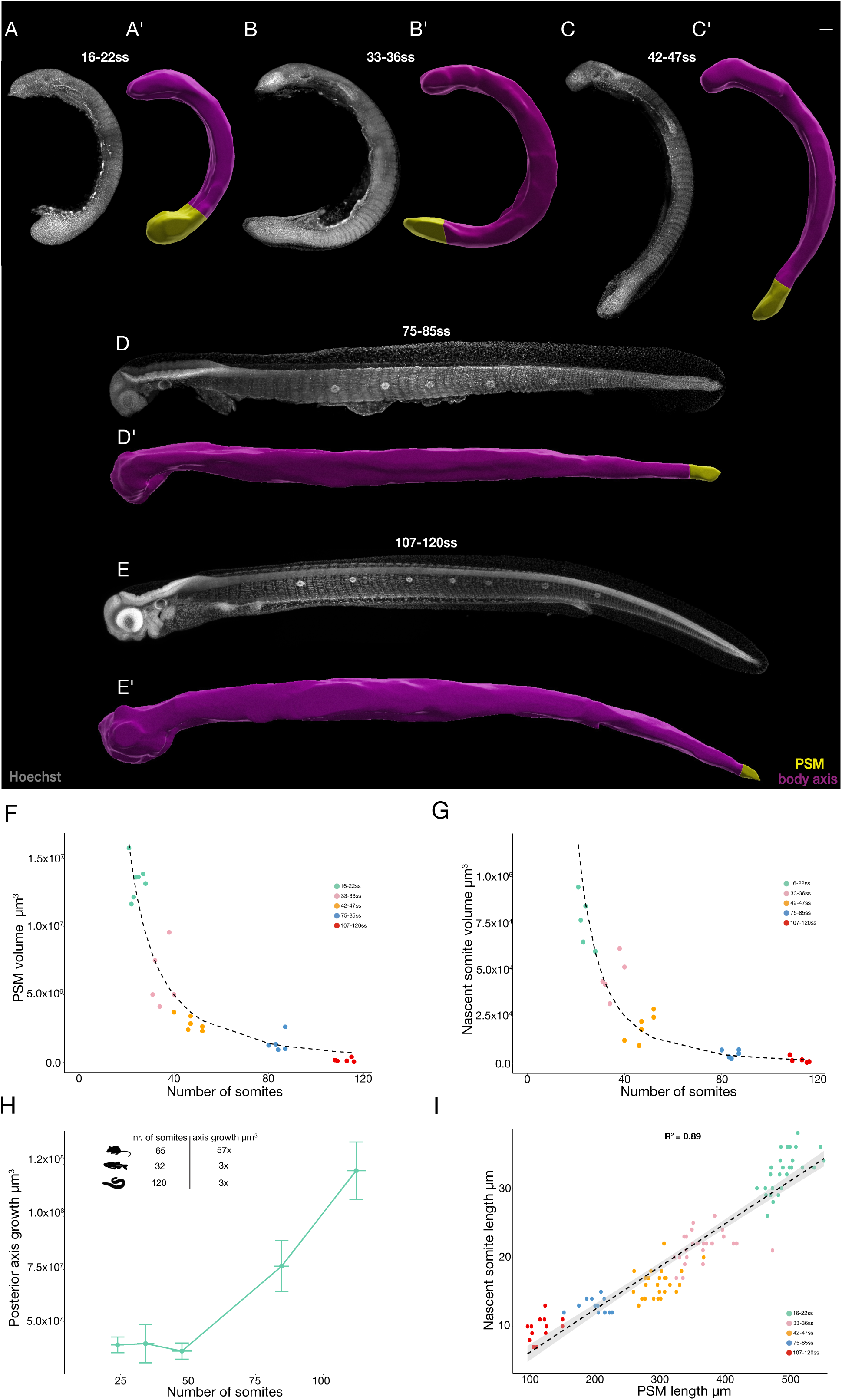
Volumetric growth and tissue scaling during axis segmentation in *A. japonica*. (A-E’) Hoechst staining of *A. japonica* fixed samples at indicated somite stages (ss) (A-E) and their corresponding 3D segmented volumes (A’-E’) yellow = unsegmented presomitic mesoderm (PSM); purple = body axis. Scale bars = 100 μm N = 9 (16–22 ss), N = 8 (33–36 ss), N = 10 (42–47 ss), N = 8 (75–85 ss), N = 9 (107–120 ss). (F) PSM volume (µm^3^) throughout somitogenesis stages 16-22ss (green N = 7), 33-36ss (pink N = 5), 42-47ss (orange N = 6), 75-85ss (blue N = 6), 107-120ss (red N = 5). (G) Nascent somite volume (µm^3^) throughout somitogenesis stages (N = 5 per stage). (H) Posterior axis volumetric growth (3.1-fold increase), from 3.9 × 10⁷ μm³ (SD ± 3.7 × 10⁶) at 20% completion to 1.2 × 10⁸ μm³ (SD ± 1.3 × 10⁷) at 94% completion. Inset shows somite number and posterior axis volumetric fold-change in *Mus musculus*, *Danio rerio* and *Anguilla japonica.* (I) Scaling of PSM to somite length (µm) throughout segmentation stages 16-22ss (green N= 27), 33-36ss (pink N= 25), 42-47ss (orange N= 26), 75-85ss (blue N= 12), 107-120ss (red N= 13). Dashed black line represents linear fit (R^2^ = 0.89), grey shading indicates the 95% confidence interval.

### Heterochronic shifts in somitogenesis using conserved clock components

To assess the temporal dynamics of axis segmentation in *A. japonica* we injected one-cell stage eel embryos with *H2A-mCherry* mRNA and recorded periodic segment addition using 4D confocal live-imaging at the 16-47 somite stages (Fig. 2, A; fig. S3A; movies S3 and S4). Our imaging data revealed an overall periodicity of somite formation of ∼25 minutes, slightly faster than that of zebrafish reared at a similar temperature (*7*). To gain a more complete picture of axis segmentation rate across all somitogenesis stages we iteratively quantified somite counts in live *A. japonica* samples over time (Fig. 2B). These data revealed an overall segmentation rate that is similar to zebrafish (*7*, *13*), but interestingly uncovered a strong stage-dependent periodicity of segment addition (Fig. 2C; fig. S3B). We observed an initial acceleration of the segmentation rate until the 33-somite stage, followed by a sharp deceleration that transitioned into a more gradual decline until the completion of somitogenesis (Fig. 2C). The rapid deceleration of the segmentation rate coincided with key developmental transitions: hatching (47-somite stage) and the switch to caudal somite formation (51-somite stage). Such non-monotonic changes in segmentation period contrast with the more gradual slowing down dynamics reported in other vertebrates (*3*, *7–9*). Next, we compared the rate of somitogenesis to overall developmental progression (Fig. 2D). This revealed specific heterochronic changes: while the onset of segmentation occurs synchronously in *A. japonica* and zebrafish, eel embryos hatch earlier in absolute time and importantly, continue to segment their body axis for at least two additional days after hatching (Fig. 2D). Hence, unlike zebrafish and medaka, where somitogenesis is restricted to mid-embryogenesis and concludes well before hatching (*19*, *20*), eels have considerably expanded the duration of the process relative to overall developmental progression. This modularity—uncoupling the termination of segmentation from other developmental milestones—enables the Japanese eel to sustain its axis segmentation program leading to a significantly increased number of segments.

**Fig. 2.**
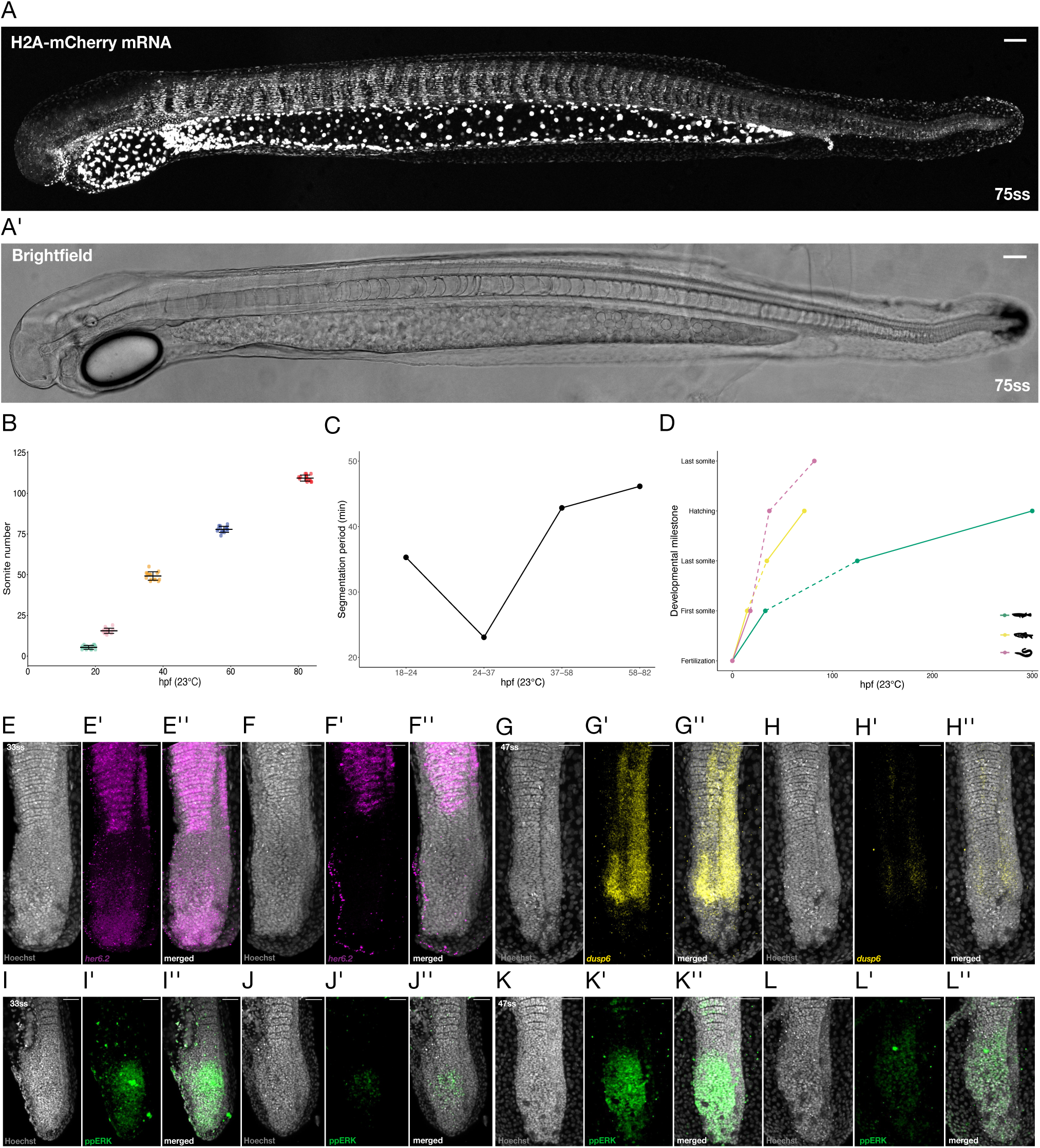
Temporally extended somitogenesis via conserved segmentation clock machinery. (A and A’) Confocal imaging of a 75ss eel live-sample injected with H2A-mcherry mRNA (grey) and corresponding brightfield image (A’) Scale bars = 100 μm; N = 3. (B) Somite counting over time in embryos raised at 23°C (N = 15 per timepoint). Data represent mean (solid line) ± SD (whiskers) hpf, hours post-fertilization. (C) Average segmentation period (min) at 23°C derived from (B): 18–24 hpf (35 min), 24–37 hpf (23 min), 37–58 hpf (43 min), and 58–82 hpf (46 min). (D) Comparative developmental timing at 23°C in *Oryzias latipes* (green), *Danio rerio* (yellow) and *Anguilla japonica* (pink). Dashed lines indicate somitogenesis stages; note heterochronic extension in *Anguilla japonica* beyond hatching stage. (E-F’’) Dynamic expression of *her6.2* (magenta) mRNA in 33-ss embryos, nuclear Hoechst staining (grey). One embryo shows high *her6.2* expression in the PSM (E’), another low (F’); N = 15 embryos across 4 stages. Scale bars = 50 μm. (G-H’’) Dynamic expression of *dusp6* (yellow) in 47-ss eel embryos. Nuclear Hoechst staining (grey). One embryo shows high *dusp6* expression in the PSM (G’), another low (H’) N = 18 embryos across 4 stages Scale bars = 50 μm. (I-L’’) Immunostaining of diphosphorylated ERK (ppERK, green) in 33-ss embryos (I to J’’, N = 4) and 47-ss embryos (K to L’’, N = 7). High and low expression phases are shown. Scale bars = 50 μm.

To determine the molecular basis of the eel segmentation clock, we used our improved *A. japonica* genome assembly to perform Hybridization Chain Reaction (HCR) (*21*) on conserved clock components across somitogenesis stages (*22*). *her6.2* showed evidence of oscillatory mRNA expression in the eel PSM (Fig. 2E to F); interestingly, for *her7* and *her1*—the core clock genes in medaka and zebrafish (*9*, *22*)— we did not detect oscillatory expression (fig. S3C to K). This suggests a degree of functional evolution in *her* paralog usage among teleosts. However, other molecular clock components remain conserved (*23–25*): we detected evidence for dynamic expression of *dusp6* (but not its paralog *dusp4*) at the transcript level (Fig. 2G to H; fig. S4A to L) and oscillatory diphosphorylated ERK (ppERK) protein levels (Fig. 2I to L) across somitogenesis stages in *A. japonica* PSMs. Notably, our HCR and immunostaining analyses did not reveal clear evidence for travelling wave patterns of oscillatory genes, as is reported in other vertebrates (*3–5*). Instead, our results are consistent with more spatially uniform ‘pulse-like’ dynamics across the PSM. Whether these ‘pulse-like’ expression dynamics are mechanistically linked to the extreme segment scaling property we report remains to be functionally tested. However, these findings raise the question about the organization of known signaling gradients in the PSM—another key parameter that impacts segment formation. Together, our data demonstrate that heterochronic shifts in the duration of axis segmentation are not dependent on changes in the core clock machinery or rate, but rather on a decoupling of somitogenesis from overall developmental progression.

### Switch of molecular PSM patterning from pre- to post-hatching somite formation

To determine the spatio-temporal expression of PSM signaling gradients and patterning genes across the extended somite formation process, we performed HCR on conserved members of the Wnt, Fgf, RA, and Notch signaling pathways in pre- and post-hatching stages. We first analyzed *msgn1* and *mespa*, conserved PSM and anterior segment markers, respectively (*5, 26*). Both markers exhibited the expected expression profiles, with *mespa* showing 2-3 stripes in the anterior PSM, while *msgn1* was expressed up to ∼70% of PSM length (Fig. 3A; fig. S5A). Interestingly, *msgn1* expression was excluded from the tailbud region until hatching (47-somite stage), after which the entire posterior PSM, including the tailbud, exhibited robust *msgn1* expression. These results indicate a spatial remodeling of molecular patterning domains in the PSM between pre-hatching and post-hatching stages. To determine whether this shift represents a broader reorganization of expression profiles in the PSM, we performed a quantitative analysis of mRNA expression patterns of conserved PSM genes across somitogenesis stages. Our results indicate that the majority of tested genes exhibited a domain change between the pre- and post-hatching stages (fig. S3, C to H; fig. S4, A to F; fig. S5, C to H; fig. S6, A to L). For instance, *lfng*—a core oscillating target of Notch signaling in mice and snakes (*4, 5*)—displayed a non-oscillatory, horseshoe-shaped domain in the PSM that persisted until the 47-somite stage, after which expression was expanded across the entire PSM (Fig. 3B; fig. S5, B to G; movie S5). Similarly, *cyp26a1* and *axin2* showed posterior PSM expression until the 47-somite stage but shifted to the anterior PSM, excluding the tailbud, by the 75-somite stage in post-hatch larvae (fig. S3, C to H; fig. S6, G to L). Intriguingly, *fgf8a* also displayed a horseshoe-shaped domain in the PSM (Fig. 3C). 3D reconstruction revealed dynamic remodeling of this domain: the posterior expression retracts over time, such that by the 75-somite stage, only an anterior domain of *fgf8a* remains in the unsegmented PSM (Fig. 3C; fig. S6, A to F; movie S6).

**Fig. 3.**
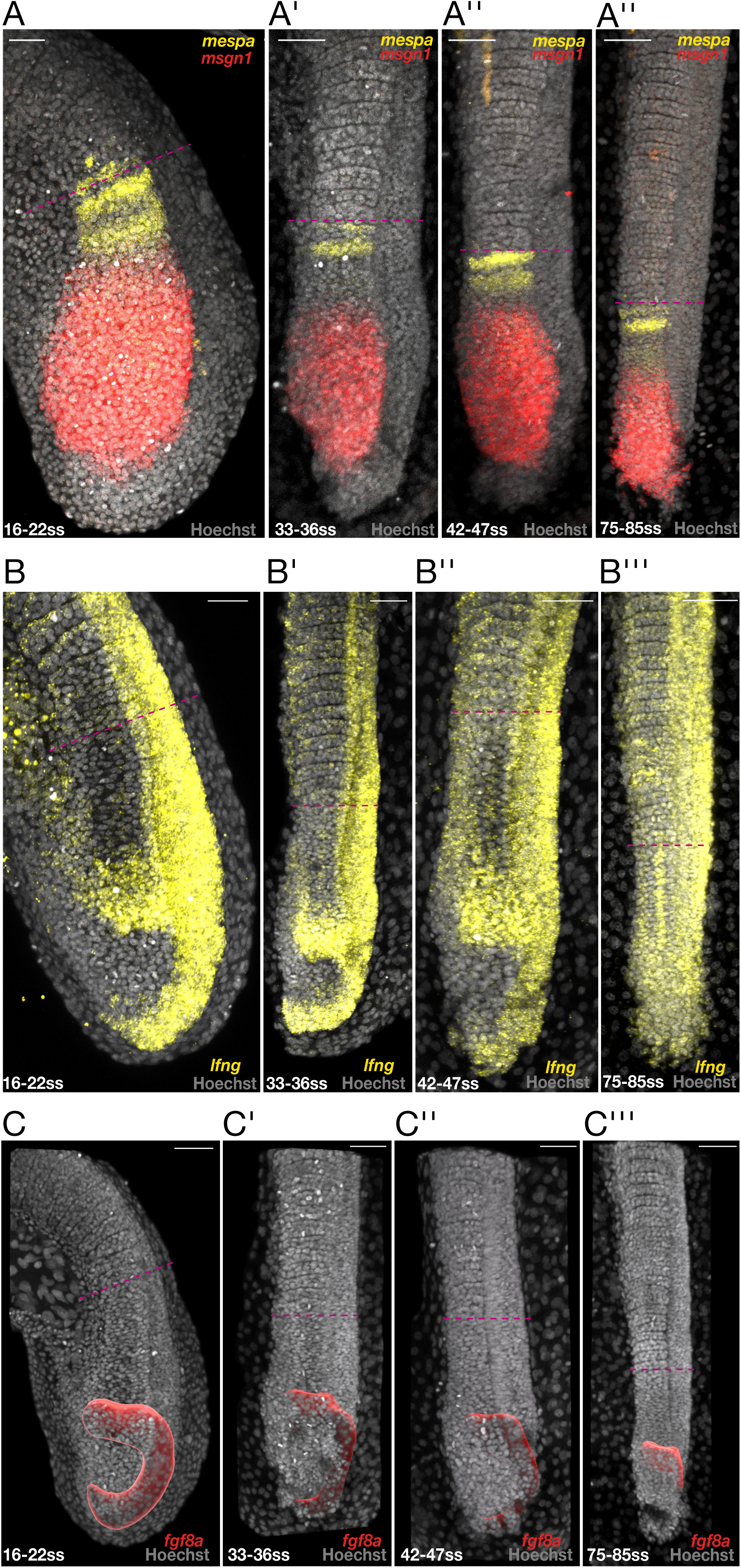
Spatio-temporal expression of PSM patterning genes in *A. japonica*. (A-A’’’’) HCR on *msgn1* (red) and *mespa* (yellow) mRNA. Hoechst = nuclear staining (grey). Dashed magenta line indicates the boundary of the last formed somite. Stages: 16-22ss (A, N = 6), 33-36ss (A’, N = 4), 42-47ss (A’’, N = 5), and 75-85ss (A’’’, N = 2). Scale bars = 50 µm. (B-B’’’) HCR on *lfng* mRNA (yellow) Hoechst = nuclear staining (grey). Dashed magenta line indicates the boundary of the last formed somite. 16-22ss (B, N = 2), 33-36ss (B’, N = 2), 42-47ss (B’’, N = 3), and 75-85ss (B’’’, N = 3). Scale bars = 50 µm (C-C’’’) 3D reconstruction of *fgf8a* mRNA expression (red) Hoechst = nuclear staining (grey) at 16-22ss (C, N = 2), 33-36ss (C’, N = 2), 42-47ss (C’’, N = 2), and 75-85ss (C’’’, N = 3). Scale bars = 70 μm.

Taken together, these data reveal unique spatiotemporal features of PSM patterning in *A. japonica*, most notably, a switch in gene expression patterns between pre- and post-hatching stages. These changes coincide with key developmental transitions: hatching and the switch to caudal segment formation. Despite this apparent modular organization of axis segmentation with each module (*i.e.* pre- and post-hatch) showing distinct expression patterns of signaling gradients in the PSM, we report a *continuous* morphological segment scaling process across all stages (Fig. 1I). Our results are therefore inconsistent with the morphological scaling process relying on simple scaling of gene expression domains in the PSM (*27*). Given the unique horseshoe-shaped expression domains we report for *fgf8a* and *lfng*, we decided to investigate the link between this spatial configuration and the organization of neuromesodermal progenitors (NMPs) in the tail.

### Expanded stemness in *time* and *space* drives extreme axis segmentation

Neuromesodermal progenitors (NMPs) are a bipotent cell population located in the posterior tail in vertebrates and are characterized by the co-expression of *sox2* and *brachyury* (*28*). To identify this population in the Japanese eel, we performed HCR on both genes followed by 3D colocalization analysis (Fig. 4A; fig. S7, A to F). While *brachyury/sox2* identifies *potential* NMPs, additional combinations of marker gene expression is used to refine their identification (*28, 29*). Using the expression of *fgf8* as a core NMP marker gene, our analysis revealed the existence of a unique horseshoe-shaped *brachyury/sox2* and *fgf8* positive domain that we interpret as reflecting *bona fide* NMP pools. Based on these expression domains, we conclude that in contrast to all previously studied vertebrates, the Japanese eel contains two NMP pools: a posterior pool in the tailbud and a more anterior pool extending into the PSM. The spatiotemporal dynamics of these pools are distinct (Fig. 4A; movie S7) by the 75-somite stage, the posterior tailbud is devoid of *sox2*/*brachyury* double-positive cells, and only the anterior NMPs remain (Fig. 4A’’’; movie S7). To functionally characterize the potential of these NMP pools, we performed microsurgical amputations at the 47-somite stage (onset of hatching). Amputating the entire PSM (and hence both NMP pools) resulted in axis truncation with no regeneration (Fig. 4, B and C; fig. S7, G and H). However, selective amputation of the posterior tailbud (by removing the posterior ⅓ of the PSM including the posterior NMP pool) did not arrest somitogenesis. Instead, these embryos continued segmentation, and by 16 hours post amputation had only slightly reduced segment numbers compared to controls (Fig. 4, B and D; fig. S7, I to L; fig. S8, A). Importantly, by 34 hours post-amputation, tailbud amputated embryos formed the full complement of somites as their control counterparts (fig. S8, B to D). These results demonstrate that after the 47-somite stage, the rostral PSM fragment containing the anterior NMP pool, is sufficient to support continued axis segmentation, independent of the tailbud proper. This finding is in contrast to data from other vertebrates, in which removal of the posterior tail-bud leads to a cessation of segmentation and axis truncation (*30, 31*).

**Fig. 4.**
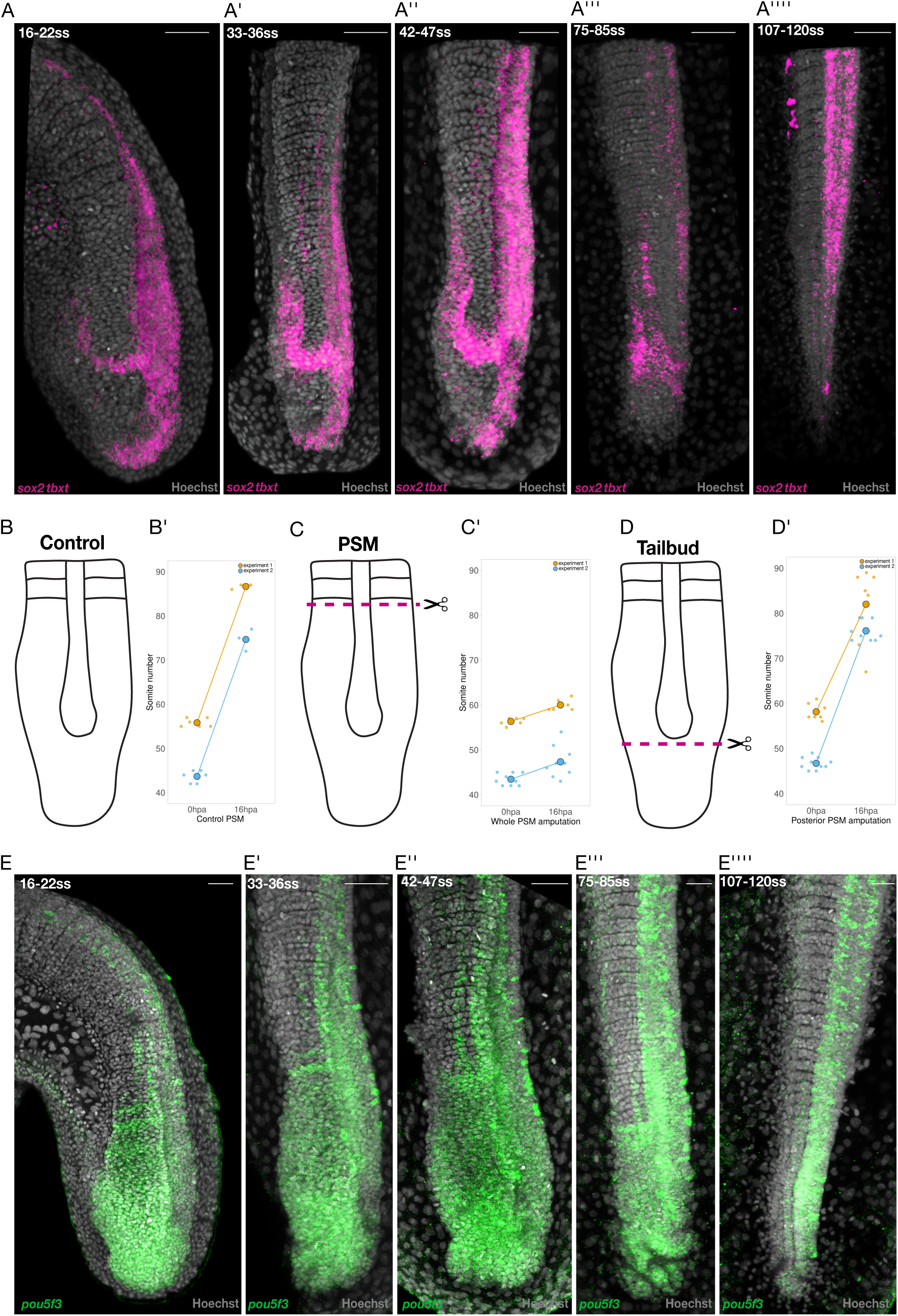
Temporally and spatially expanded stemness drives extended axis segmentation. (A-A’’’’) 3D colocalised signal of *sox2* and *tbxt* mRNA expression (magenta). Hoechst = nuclear staining (grey). 16-22ss (A, N = 2), 33-36ss (A’, N = 2) Scale bars = 70 µm, 42-47ss (A’’, N = 3), and 75-85ss (A’’’, N = 3) 107-120ss (A’’’’, N = 3). Scale bars = 50 μm. (B-D’) Schematics and results of PSM amputation experiments at ∼ 47-ss. hpa, hours post-amputation; temperature, 26°C, SD = standard deviation. Control non-amputated (B-B’) experiment 1 (orange dots) control somites at 0hpa 55.83 (SD ± 0.98) N= 6, control somites at 16hpa 86.67 (SD ± 0.58) N= 3, experiment 2 (blue dots) control somites at 0hpa 43.67 (SD ± 1.37) N= 6, number of somites at 16hpa 74.67 (SD ± 2.52) N = 3. Whole PSM amputation (C-C’) magenta dotted line shows the amputation plane, experiment 1 (orange dots) number of somites at 0hpa 56.33 (SD ± 0.82) N= 6, number of somites at 16hpa 60 (SD ± 1.26) N= 6, experiment 2 (blue dots) at 0hpa 43.44 (SD ± 1.33) N= 9, number of somites at 16hpa 47.33 (SD ± 3.5) N= 9. Tailbud (posterior PSM) amputation (D-D’) magenta dotted line shows the amputation plane, experiment 1 (orange dots) number of somites at 0hpa 58.14 (SD ± 1.86) N= 7, number of somites at 16hpa 82 (SD ± 8.56) N= 7, experiment 2 (blue dots) number of somites at 0hpa 46.7 (SD ± 1.34) N = 10, tailbud (posterior PSM) amputation number of somites at 16hpa 76.1 (SD ± 2.13) N = 10. (E-E’’’’) HCR on *pou5f3* (*oct4*) mRNA (green) Hoechst = nuclear staining (grey) at 16-22ss (E, N = 4), 33-36ss (E’, N = 3), 42-47ss (E’’, N = 2), Scale bars = 50 μm and 75-85ss (E’’’, N = 3) 107-ss (E’’’’, N = 2). Scale bars = 30 μm.

We next investigated how *A. japonica* NMPs maintain their potential for an extended duration (over two days post-hatching). In snakes, the heterochronic extension of *Oct4* expression in the tail was identified as the basis for the sustained stemness program that underlies axis expansion (*32*). HCR analysis of *pou5f3* (the teleost *Oct4* ortholog) in *A. japonica* revealed expression throughout the PSM that persisted until the 107-somite stage (Fig. 4E; fig. S8, E to J). This stands in contrast to *pou5f3/Oct4* dynamics in medaka, zebrafish and mouse, where expression is temporally restricted and localized to the posterior tail (*32–34*). Our data suggests that the maintenance of Pou5-family pluripotency factors and their broader expression domain in the tail are a convergent evolutionary strategy shared between eels and snakes to support prolonged axis segmentation. Consistent with this, we also observed sustained expression of the heterochronic stemness regulators *lin28a/b* (*35–37*) throughout *A. japonica* somitogenesis (fig. S9, A to F). Since *Hox13* cluster activation is known to terminate the segmentation program by repressing Lin28 and WNT/FGF signaling leading to the exhaustion of tailbud progenitors in amniotes (*32*, *35*), we examined *hoxb13a* expression in *A. japonica.* We found that *hoxb13a* remains undetectable in the PSM until the 107-somite stage (fig. S9, G to H)—coinciding with the final downregulation of *pou5f3*. Our data suggests that *Hox13* activation in the Japanese eel is temporally delayed and that its molecular function more closely aligns with its reported role in amniotes (*32*, *35*). This contrasts with *Hox13* cluster expression dynamics and function in zebrafish where it is active during early somitogenesis and supports the maintenance of axial progenitors (*38*). Taken together, our results indicate that delayed activation of the *Hox13* cluster and the temporal extension of *pou5f3* expression (*32*), maintain the extended stemness program in the *A. japonica* PSM.

In this study, we identify the temporal extension and spatial expansion of *Oct4* expression in the tail as a convergent evolutionary signature shared between snakes and eels. This demonstrates that evolution targets common regulatory nodes—such as the pluripotency network—to drive prolonged axis segmentation across vast phylogenetic distances (*39*, *40*). Beyond this shared strategy, we reveal tinkering at the level of gene expression and NMP domains that can explain the prolonged segmentation program in the Japanese eel. Furthermore, we describe a scaling regime that sustains expanded segmentation in the absence of significant axis growth. Collectively, our results demonstrate how developmental modularity—the heterochronic shift of segmentation into post-hatching stages and the tunability of axis scaling—enables the diversification of the vertebrate body plan. By identifying molecular and cellular signatures of the heterochronic shift underlying an extreme case of axis segmentation in vertebrates, our findings contribute to a deeper understanding of the heterochronic processes first described in classical work (*41*, *42*). Given that eel-shaped body plans have evolved multiple times independently in vertebrates (*40*), this study provides a foundation for comparative approaches to determine whether evolution achieved these convergent shapes by taking similar or divergent developmental trajectories.

## Acknowledgments

We would like to thank all members of the Aulehla lab for input and feedback on the project. We would also like to thank Cantas Alev and members of his lab for comments and feedback during the project. We would like thank Nicoletta Petridou and members of her lab for providing mRNA aliquots for injections. We would like to extend our warmest thanks and appreciation to all the staff at Minamiizu Field Station, Fisheries Technology Institute, Japan Fisheries Research and Education Agency and at National Institute for Basic Biology, Nishigonaka 38, Myodaiji, Okazaki NIBB for their hospitality, help and support.

## Funding

The European Molecular Biology Laboratory (EMBL)

European Research Council Consolidator Grant ERC-CoG-866537 (A.A.)

European Molecular Biology Laboratory core funding (A.S., A.A.)

EMBL Planetary Biology seed grant (A.S.)

NIBB collaborative research program 25NIBB348 (A.S.)

German Research Foundation HI2214/1-1, Project number: 530763738 (L.H.)

## Author contributions

Conceptualization: AS, KNa, YK, AA

Methodology: AS, LH, MH, YK, AA

Validation: AS, MH

Formal analysis: AS, KNo, LH

Investigation: AS, KNo

Resources: MH, KNa, YK, AA

Data Curation: AS, LH, KNo

Writing - original draft: AS, AA

Writing - review and editing: AS, KNo, LH, MH, KNa, YK

Visualization: AS

Supervision: KNa, AA

Project administration: YK, KNa, AA

Funding acquisition: AS, AA

## Data, code, and materials availability

The *A. japonica* draft genome assembly is available at DDBJ (accession nos. BAAJLN010000001-BAAJLN010000107). All cDNA for HCR probe generation is provided in Data S1. All other data are available in the main text or supplementary materials. All imaging and source data are deposited in BioImage Archive, any materials used in this study are available upon request from the corresponding author.

Supplementary Materials

Materials and Methods

Figs. S1 to S9

References (43–58)

Movies S1 to S7

Data S1

## Materials and Methods

### *A. japonica* husbandry, animal ethics and IVF induction

*A. japonica* eels were maintained in a system with a flow rate of 1-2L/ min, using seawater at 15-20°C at the Minamiizu Field Station, Fisheries Technology Institute (FTI), Japan Fisheries Research and Education Agency (FRA). Animal experiments were performed under approval by the Institute Animal Care and Use Committee of the Fisheries Technology Institute (permission code: 23016). Eel larvae were obtained by *in vitro* artificial fertilization (IVF). Detailed protocols of assisted reproduction in Japanese eels are described elsewhere (*10*, *43*). In brief, maturation of male eels was induced by administration of recombinant Japanese eel luteinizing hormone, and milt was collected (*43*) and cryopreserved in liquid nitrogen until IVF (*44*). Female eels received intraperitoneal injections of recombinant Japanese eel follicle-stimulating hormone until they reached full maturity. Thereafter, in order to induce the final oocyte maturation and ovulation, the female eels were administered with a combination of an analogue of luteinizing hormone-releasing hormone and pimozide, followed by subsequent injection of 17α-hydroxyprogesterone (*10*). After confirmation of ovulation, eggs were gently stripped from the female eels and immediately inseminated with the cryopreserved sperm samples which have been thawed prior to use and diluted with an artificial seminal plasma (*45*).

### mRNA injections in *A. japonica* eggs

Freshly fertilized eel embryos were placed in *1⁄3* seawater (*2/3* freshwater) to prevent floating. An *Oryzias latipes* injection mould (1.1mm wide) was used to prepare an agarose coated injection plate. 0.6% agarose in H_2_O was used to prevent embryo bursting and a blunt forceps was used to orient the embryos in the wells. Modified zebrafish injection needles (wider needle opening) were used to enable penetration of the eel embryo chorion while ensuring sufficient injection volume. Injections targeted both the yolk and the developing single-cell. H2B-GFP or H2A-mcherry mRNA were injected with a final concentration of 15 ng/µl. Immediately after the injection procedure eel embryos were transferred to seawater and kept at 23-24°C.

### Hybridization chain reaction (HCR) and immuno-histochemistry

*A. japonica* eel embryos were staged at 16ss (24hpf), 33ss (33hpf), 47ss (37hpf), 75ss (58hpf), 107ss (82hpf) then fixed in 4% PFA in PtW for 24 hours at 4°C. Embryos were subsequently washed 3x in PtW followed by dehydration and storage in MeOH at -20°C. All hybridization chain reaction (HCR) (*46*) probes were designed in-silico (*47*) and ordered as pooled DNA oligos from Integrated DNA technologies (IDT). A full list of all targeted genes and their corresponding cDNA sequences are provided in Supplementary Data 1. The homologous eel genes and cDNAs were identified by performing BLASTN (v2.2.30) searches against an in-house *A. japonica* draft genome assembly (DDBJ accession nos. BAAJLN010000001-BAAJLN010000107) using an E-value cutoff of 1e-5. cDNA sequences of *Mus musculus*, *Oryzias latipes*, and *Danio rerio* retrieved from the *Ensembl* database were used as queries. This approach was complemented by using *TOGA v.1.0* (Tool to infer Orthologs from Genome Alignments) (*48*) to annotate protein coding genes in the genome of the Japanese eel *Anguilla japonica* (GCA_916313395.1). As a reference we chose the Pacific tarpon *Megalops cyprinoides* (GCF_013368585.1). We aligned the Japanese eel genome to the Pacific tarpon genome using a previously-established pipeline (*49*). To account for the large phylogenetic distance between reference and query we ran LASTZ v.1.04.15 with increased sensitivity (K= 2200, L= 2800, Y= 9400, H= 2000, and the LASTZ default scoring matrix) (*50*). To generate and chain local alignments, we used axtChain (*51*) with default parameters except for linearGap= loose. We used RepeatFiller (*52*) with default parameters to add missed repeat-overlapping local alignments to the alignment chains and chainCleaner (*53*) with default parameters except for minBrokenChainScore= 75,000 and -doPairs to improve alignment specificity. Finally, we ran TOGA to annotate genes, classify gene intactness and extract gene sequences for subsequent hybridization chain reaction (HCR) and immuno-histochemistry. Our data confirmed the cDNA for all assessed genes in *A. japonica*, additionally it indicated that *cyp26a1* is likely a fusion transcript with *cyp26b1*-like and that *mespa* is likely a fusion transcript with *mespb*, our HCR probes were designed to cover the entire gene in both cases. All buffers and reagents were ordered from Molecular instruments (MI). We followed the publicly available HCR protocol (*46*), the following HCR amplifiers were used B1-546, B2-546, B2-647 (Molecular Instruments, MI). Hoechst 33342 (Thermo Fisher #H3570) was used with a dilution of 1:500 of 10 mg/ml stock solution as a nuclear label. For immunostainings; fixed embryos were stored in 100% Methanol and rehydrated with a series of graded PBST (75%MeOH, 50%MeOH, 25% MeOH). The samples were permeabilized by placing in detergent solution (*46*) for 30 minutes. Samples were then blocked for 2 hours at room temperature (blocking solution 2% BSA, 5% NGS in PBST). Samples were then incubated in the primary antibodies: anti-MAP Kinase (diphosphorylated ERK1,2) antibody Mouse monoclonal (Sigma-Aldrich M9692) and anti-PH3 antibody rabbit monoclonal (Cell signaling technology #3642) at 1:200 overnight at 4°C. Embryos were then washed 3x 1 hour in PBST followed by secondary antibodies incubation: Goat anti-Rabbit 568 Alexa Fluor 568 (Thermo Fisher #A-11011) and Goat anti-Mouse Alexa Fluor 647 (Thermo Fisher #A21235) and Hoechst 33342 (Thermo Fisher #H3570) 1:500 overnight at 4°C. After washing 3x 1 hour in PBST embryos were mounted in 8-well glass-bottomed dishes (Lab-Tek Chambered #1 Borosilicate Coverglass System 155411, T.S) using low melting agarose (0.6 to 1%) (Biozyme Plaque Agarose #840101) and imaged.

### Live-imaging sample preparation and microscopy

For confocal 4D live-imaging eel embryos were manually dechorionated using sharp forceps. 1x Tricaine (Sigma-Aldrich #A5040-25G) was used to anaesthetize dechorionated eel embryos (20 mg/ml – 20x stock solution diluted in seawater). Anesthetized embryos were then mounted using low melting agarose (0.6 to 1%) (Biozyme Plaque Agarose #840101). We then proceeded to carefully remove the solidified agarose from the tail section as previously described in zebrafish (*54*). Imaging was performed using 8-well glass-bottomed dishes (Lab-Tek Chambered #1 Borosilicate Coverglass System 155411, T.S). Laser-scanning confocal Leica SP5 and Leica Stellaris microscopes with 20x and 40x objectives were used during image acquisition for both 4D live imaging and HCR stained embryos. Embryos were imaged at the 16ss, 33ss, 47ss, 75ss and 107ss. For live imaging experiments; temperature was set at 23°C, 24°C or 26°C and was measured throughout imaging using a temperature sensor (SHT4x, Sensirion). For tail amputations hatched embryos at stage 42-47ss were briefly anesthetized in 1x Tricaine in seawater, then using two sharp forceps and a blade either the whole PSM (up to the last formed somite) was cut or the last third of the PSM (directly below the end of the notochord), control embryos were placed in 1x Tricaine but no cut was performed. Directly after tail cutting embryos were recovered in seawater. Subsequent counting of somite number was done manually at 16 hours post amputation for all embryos and an end point of 34 hours post amputation for a subset of embryos, representative images before and after amputation were taken on a NIKON stereoscope (SMZ18-DBL-1) equipped with digital camera (DS-Ri2).

### Data analysis

Open-source ImageJ/Fiji software (*55*) was used for image analysis of all microscopy imaging. Image stitching was performed using 2D and 3D plug-ins on ImageJ/Fiji. For all HCR quantifications a line (width 80 pixels) was drawn on ImageJ/Fiji from the posterior tip of the PSM until the first somite boundary (defined by Hoechst staining). Absolute (μm) and relative (percentage of PSM) distances were then extracted for each trace across all developmental stages assayed. Normalization by dividing with the largest value for each trace was used to calculate the relative distances. For immuno-histochemistry and mitotic index analysis both PH3*+* cells and all Hoechst stained PSM cells were segmented in 3D using *Imaris* software and the number of cells was quantified in each somitogenesis stage. The colocalization of *sox2* and *tbxt* expression in 3D was done using the *COLOC* module in *Imaris*. Data were plotted using *ggplot2* and *gganimate* in R software or using *PlotTwist* (*56*) and *PlotsofData*, Super*PlotsofData* (*57*). For all volumetric analysis *Imaris* was used to create 3D surfaces and to quantify the volume of PSM, somites and segmented body axis across different somitogenesis stages. Somitogenesis data on mouse, zebrafish and medaka were obtained from published reports(*5*, *7–9*, *14*, *58*). The mitotic index (MI) was calculated as the percentage of PH3*+* cells from the total number of cells in the unsegmented tail region. *Imaris* was used to segment and count nuclei from the unsegmented tail region across all somitogenesis stages reported. All statistical analysis including: Pearson’s product moment correlations and Welch two sample t-tests were calculated and plotted in R software version 4.2.2.

## Supplementary Figure Legends

Fig. S1. PSM and nascent somite quantifications.

(A) 16-22ss embryo Hoechst = nuclear staining (grey), magenta dotted line = PSM length (µm) yellow dotted line = nascent somite length (µm) scale bar = 50 µm N= 27 (A’) 33-36ss Hoechst = nuclear staining (grey) magenta dotted line = PSM length (µm) yellow dotted line = nascent somite length (µm) scale bar = 50 µm N= 25 (A’’) 42-47ss Hoechst = nuclear staining (grey) magenta dotted line = PSM length (µm) yellow dotted line = nascent somite length (µm) scale bar = 50 µm N= 26 (A’’’) 75-85ss Hoechst = nuclear staining (grey) magenta dotted line = PSM length (µm) yellow dotted line = nascent somite length (µm) scale bar = 50 µm N= 12 (A’’’’) 107-120ss Hoechst = nuclear staining (grey) magenta dotted line= PSM length (µm) yellow dotted line = nascent somite length (µm) scale bar = 50 µm N= 13 (B) PSM length (µm) throughout somitogenesis 16ss N= 27, 33ss N= 25, 47ss N= 26, 75ss N= 12, 107ss N= 13 green dots = mean, green whiskers = ± SD (standard deviation) (C) Nascent somite length µm throughout somitogenesis 16ss N= 27, 33ss N= 25, 47ss N= 26, 75ss N= 12, 107ss N= 13 yellow dots = mean, yellow whiskers = ± SD (standard deviation) (D) limited posterior axis elongation throughout somitogenesis (3-fold) from 1577 µm (SD ± 104) at 20% somitogenesis completion to 4728 µm (SD ± 358) at 94% somitogenesis completion (E) limited posterior axis area growth over somitogenesis (4-fold) from 244669µm^2^ (SD ± 35647) at 20% somitogenesis completion to 978645 µm^2^ (SD ± 139394) at 94% somitogenesis completion (F) mitotic index (MI) measured as the percentage of PH3+ cells in 3D unsegmented tail region across different somitogenesis stages 16-22ss (green) 6.21% (SD ± 0.42) N= 5, 33-36ss (pink) 8.25% (SD ± 1.34) N= 4, 42-47ss (orange) 2.95% (SD ± 1.78) N= 7, 75-85ss (blue) 2.22% (SD ± 1.1) N= 3, 107-120ss (red) 0.46% (SD ± 0.34) N= 4 (G) PH3+ immunostaining (green) on a 16-22ss eel embryo nuclei = Hoechst (grey) scale bar = 50 µm N= 5 (G’) PH3+ immunostaining (green) on a 33-36ss eel embryo nuclei= Hoechst (grey) N= 4 scale bar= 30 µm (G’’) PH3+ immunostaining (green) on a 42-47ss eel embryo nuclei = Hoechst (grey) N= 7 scale bar = 30 µm (G’’’) PH3+ immunostaining (green) on a 75-85ss eel larvae nuclei = Hoechst (grey) N= 3 scale bar = 30 µm (G’’’’) PH3+ immunostaining (green) on a 107-120ss eel larvae nuclei= Hoechst (grey) N= 4 scale bar = 20 µm (H) segmented body axis of a 107-120ss eel Hoechst = nuclear staining (black) scale bar = 300 µm (I) Total number of somites formed in *Oryzias latipes* 35, *Danio rerio* 32 and *Anguilla japonica* 120 (J) Total number of abdominal somites formed in *Oryzias latipes* 15, *Danio rerio* 17 and *Anguilla japonica* 51 (K) Total number of caudal somites formed in *Oryzias latipes* 20, *Danio rerio* 15 and *Anguilla japonica* 69.

Fig. S2. Abdominal and caudal somites and *hoxd12a* expression boundaries

(A) ratio of abdominal somites to total somite count in *Oryzias latipes* 0.43, *Danio rerio* 0.53 and *Anguilla japonica* 0.43 (B) ratio of caudal somites to total somite count in *Oryzias latipes* 0.57, *Danio rerio* 0.47 and *Anguilla japonica* 0.57 (C) 75-85 somite stage eel embryo nuclei= Hoechst (grey) N= 4 scale bar = 100 µm red arrow = position of anus (C’) 75-85 somite stage eel embryo *hoxd12a* mRNA (yellow) red dotted line = anterior boundary of *hoxd12a* expression in the neural tube N= 4 scale bar = 100 µm (C’’) 75-85 somite stage eel embryo *hoxd12a* mRNA (yellow) Hoechst = nuclei (grey) red dotted line = anterior boundary of *hoxd12a* expression N= 4 scale bar = 100 µm (D) 107-120 somite stage eel Hoechst = nuclei (grey) red arrow = position of anus N= 4 scale bar =100 µm (D’) 107-120 somite stage eel embryo *hoxd12a* mRNA (yellow) red dotted line = anterior boundary of *hoxD12a* expression in the neural tube N= 4 scale bar = 100 µm (D’’) 107-120 somite stage eel embryo *hoxd12a* mRNA (yellow) Hoechst = nuclei (grey) red dotted line = anterior boundary of *hoxd12a* expression in the neural tube N= 4 scale bar = 100 µm.

Fig. S3. 4D-imaging and HCR on segmentation clock genes

(A) Snapshots from time-lapse confocal live-imaging of H2A-mcherry injected eel embryo at 33 somite stage scale bar =100 µm, bottom left= time in hours temperature = 26°C N= 3 (B) Quantification of segmentation period (min) from confocal 4D live-imaging of eel embryos at different developmental stages from 16-ss = 28 min (SD ± 0.22), 33-ss = 18 min (SD ± 0.69), 47ss= 30 min (SD ± 2.31) temperature = 26°C, mean= black line, (± SD) = whiskers N= 2 at 16ss N= 3 at 33ss and N= 7 at 47ss (C-C’’’’) HCR on *her7* mRNA (yellow) and *cyp26a1* mRNA (blue) Hoechst = nuclear staining (grey) at the 16-22ss scale bar = 50 µm N= 2 embryos (D-D’’’’) HCR on *her7* mRNA (yellow) and *cyp26a1* mRNA (blue) Hoechst= nuclear staining (grey) at the 33-36ss scale bar = 50 µm N = 4 embryos (E-E’’’’) HCR on *her7* mRNA (yellow) and *cyp26a1* mRNA (blue) Hoechst = nuclear staining (grey) at the 42-47ss scale bar = 50 µm N= 3 embryos (F-F’’’’) HCR on *her7* mRNA (yellow) and *cyp26a1* mRNA (blue) Hoechst = nuclear staining (grey) at the 75-85ss scale bar = 30 µm N = 2 embryos (G-G’’’’) HCR on *her7* mRNA (yellow) and *cyp26a1* mRNA (blue) Hoechst = nuclear staining (grey) at the 107-120ss scale bar = 30 µm N= 3 embryos (H) Average expression of *her7* normalised to percentage of PSM length across 5 somitogenesis stages, 16-22ss (green) N= 2, 33-36ss (pink) N= 4, 42-47ss (orange) N= 3, 75-85ss (blue) N= 2, 107-120ss (red) N= 3 (H’) average expression of *cyp26a1* normalised to percentage of PSM length across 5 somitogenesis stages, 16-22ss (green) N= 2, 33-36ss (pink) N= 4, 42-47ss (orange) N= 3, 75-85ss (blue) N= 2, 107-120ss (red) N= 3 (I-I’’) HCR on *her1* mRNA (yellow) and Hoechst = nuclear staining (grey) at the 16-22ss scale bar = 50 µm N = 4 embryos (J-J’’) HCR on *her1* mRNA (yellow) and Hoechst = nuclear staining (grey) at the 33-36ss scale bar = 50 µm N= 3 embryos (I-I’’) HCR on *her1* mRNA (yellow) and Hoechst = nuclear staining (grey) at the 42-47ss scale bar = 50 µm N= 5 embryos.

Fig. S4. HCR on *her6.2*, *dusp4/6*, and *pcna*

(A-A’’’’) HCR on *her6.2* mRNA (magenta) and *dusp4* mRNA (green) Hoechst = nuclear staining (grey) at 16-22ss scale bar = 50 µm N= 6 embryos (B-B’’’’) HCR on *her6.2* mRNA (magenta) and *dusp4* mRNA (green) Hoechst = nuclear staining (grey) at 33-36ss, same embryo as in Figure 2D, scale bar = 50 µm N= 3 embryos (C-C’’’’) HCR on *her6.2* mRNA (magenta) and *dusp4* mRNA (green) Hoechst = nuclear staining (grey) at the 42-47ss scale bar = 50 µm N= 4 embryos (D-D’’’’) HCR on *her6.2* mRNA (magenta) and *dusp4* mRNA (green) Hoechst = nuclear staining (grey) at 75-85ss scale bar = 30 µm N = 2 embryos (E-E’’’’) HCR on *her6.2* mRNA (magenta) and *dusp4* mRNA (green) Hoechst = nuclear staining (grey) at the 107-120ss scale bar = 30 µm N= 2 (F) Average expression of *her6.2* normalised to percentage of PSM length across 5 somitogenesis stages, 16-22ss (green) N= 6, 33-36ss (pink) N= 3, 42-47ss (orange) N= 4, 75-85ss (blue) N= 2, 107-120ss (red) N= 2 (F’) Average expression of *dusp4* normalised to percentage of PSM length across 5 somitogenesis stages, 16-22ss (green) N= 6, 33-36ss (pink) N= 3, 42-47ss (orange) N= 4, 75-85ss (blue) N= 2, 107-120ss (red) N= 2 (G-G’’’’) HCR on *pcna* mRNA (red) and *dusp6* mRNA (yellow) Hoechst = nuclear staining (grey) at 16-22ss scale bar = 50 µm N= 3 embryos (H-H’’’’) HCR on *pcna* mRNA (red) and *dusp6* mRNA (yellow) Hoechst = nuclear staining (grey) at 33-36ss scale bar = 50 µm N= 6 embryos (I-I’’’’) HCR on *pcna* mRNA (red) and *dusp6* mRNA (yellow) Hoechst = nuclear staining (grey) at 42-47ss, same embryo as in Figure 2F scale bar = 30 µm N= 5 embryos (J-J’’’’) HCR on *pcna* mRNA (red) and *dusp6* mRNA (yellow) Hoechst = nuclear staining (grey) at 75-85ss scale bar = 30 µm N = 3 embryos (K-K’’’’) HCR on *pcna* mRNA (red) and *dusp6* mRNA (yellow) Hoechst = nuclear staining (grey) at 107-120ss scale bar = 30 µm N= 2 embryos (L) average expression of *pcna* normalised to percentage of PSM length across 5 somitogenesis stages, 16-22ss (green) N= 3, 33-36ss (pink) N= 6, 42-47ss (orange) N= 5, 75-85ss (blue) N= 3, 107-120ss (red) N= 2 (L’) average expression of *dusp6* normalised to percentage of PSM length across 5 somitogenesis stages, 16-22ss (green) N= 3, 33-36ss (pink) N= 6, 42-47ss (orange) N= 5, 75-85ss (blue) N= 3, 107-120ss (red) N= 2.

Fig. S5. HCR on *msgn1*, *lfng*, and *wnt3*

(A) average expression of *msgn1* across different somitogenesis stages normalised to percentage of PSM length 16-22ss (green) N= 5, 33-36ss (pink) N= 4, 42-47ss (orange) N= 5, 75-85ss (blue) N= 2, 107-120ss (red) N= 5 (B) average expression of *lfng* across different somitogenesis stages normalised to percentage of PSM length 16-22ss (green) N= 2, 33-36ss (pink) N= 2, 42-47ss (orange) N= 3, 75-85ss (blue) N= 3, 107-120ss (red) N= 3 (C-C’’’’) HCR on *lfng* mRNA (yellow) and *wnt3* mRNA (blue) Hoechst = nuclear staining (grey) at the 16-22ss scale bar = 50 µm N= 2 embryos (D-D’’’’) HCR on *lfng* mRNA (yellow) and *wnt3* mRNA (blue) Hoechst = nuclear staining (grey) at the 33-36ss bar = 50µm N= 2 (E-E’’’’) HCR on *lfng* mRNA (yellow) and *wnt3* mRNA (blue) Hoechst = nuclear staining (grey) at the 42-47ss scale bar = 50 µm N= 3 (F-F’’’’) HCR on *lfng* mRNA (yellow) and *wnt3* mRNA (blue) Hoechst = nuclear staining (grey) at the 75-85ss scale bar = 50 µm N= 3 (G-G’’’’) HCR on *lfng* mRNA (yellow) and *wnt3* mRNA (blue) Hoechst = nuclear staining (grey) at the 107-120ss scale bar = 50 µm N= 2 (H) average expression of *wnt3* across different somitogenesis stages normalised to percentage of PSM length 16-22ss (green) N= 2, 33-36ss (pink) N= 2, 42-47ss (orange) N= 3, 75-85ss (blue) N= 3, 107-120ss (red) N= 2.

Fig. S6. HCR on *fgf8a/17, axin2,* and *dll1*

(A-A’’’’) HCR on *fgf8a* mRNA (magenta) and *fgf17* mRNA (green) Hoechst = nuclear staining (grey) at the 16-22ss scale bar = 50 µm N= 2 embryos (B-B’’’’) HCR on *fgf8a* mRNA (magenta) and *fgf17* mRNA (green) Hoechst = nuclear staining (grey) at the 33-36ss scale bar = 50 µm N = 2 embryos (C-C’’’’) HCR on *fgf8a* mRNA (magenta) and *fgf17* mRNA (green) Hoechst = nuclear staining (grey) at the 42-47ss scale bar = 50 µm N = 2 embryos (D-D’’’’) HCR on *fgf8a* mRNA (magenta) and *fgf17* mRNA (green) Hoechst = nuclear staining (grey) at the 75-85ss scale bar = 50 µm N= 3 embryos (E-E’’’’) HCR on *fgf8a* mRNA (magenta) and *fgf17* mRNA (green) Hoechst = nuclear staining (grey) at the 107-120ss scale bar = 50 µm N = 3 embryos (F) average expression of *fgf8a* across different somitogenesis stages normalised to percentage of PSM length 16-22ss (green) N = 2, 33-36ss (pink) N = 2, 42-47ss (orange) N = 2, 75-85ss (blue) N = 2, 107-120ss (red) N = 3 (F’) average expression of *fgf17* across different somitogenesis stages normalised to percentage of PSM length, 16-22ss (green) N = 2, 33-36ss (pink) N = 2, 42-47ss (orange) N = 2, 75-85ss (blue) N = 3, 107-120ss (red) N= 3 (G-G’’’’) HCR on *dll1* mRNA (red) and *axin2* mRNA (cyan) Hoechst = nuclear staining (grey) at the 16-22ss scale bar = 50 µm N= 3 embryos (H-H’’’’) HCR on *dll1* mRNA (red) and *axin2* mRNA (cyan) Hoechst = nuclear staining (grey) at the 33-36ss scale bar = 50 µm N= 4 embryos (I-I’’’’) HCR on *dll1* mRNA (red) and *axin2* mRNA (cyan) Hoechst = nuclear staining (grey) at the 42-47ss scale bar = 50 µm N= 2 embryos (J-J’’’’) HCR on *dll1* mRNA (red) and *axin2* mRNA (cyan) Hoechst = nuclear staining (grey) at the 75-85ss scale bar = 50 µm N= 2 embryos (K-K’’’’) HCR on *dll1* mRNA (red) and *axin2* mRNA (cyan) Hoechst = nuclear staining (grey) at the 107-120ss scale bar = 50 µm N= 3 embryos (L) average expression of *dll1* across different somitogenesis stages normalised to percentage of PSM length 16-22ss (green) N = 3, 33-36ss (pink) N = 4, 42-47ss (orange) N= 2, 75-85ss (blue) N = 2, 107-120ss (red) N = 3 (L’) average expression of *axin2* across different somitogenesis stages normalised to percentage of PSM length 16-22ss (green) N= 3, 33-36ss (pink) N= 4, 42-47ss (orange) N= 2, 75-85ss (blue) N= 2, 107-120ss (red) N= 3.

Fig. S7. Axial progenitors and microsurgical amputation

(A-A’’’’) HCR on *tbxt* mRNA (cyan) and *sox2* mRNA (red) Hoechst = nuclear staining (grey) at the 16-22ss scale bar = 50 µm N = 2 embryos (B-B’’’’) HCR on *tbxt* mRNA (cyan) and *sox2* mRNA (red) Hoechst = nuclear staining (grey) at the 33-36ss scale bar = 50 µm N = 2 embryos (C-C’’’’) HCR on *tbxt* mRNA (cyan) and *sox2* mRNA (red) Hoechst = nuclear staining (grey) at the 42-47ss scale bar = 30 µm N = 3 embryos (D-D’’’’) HCR on *tbxt* mRNA (cyan) and *sox2* mRNA (red) Hoechst = nuclear staining (grey) at the 75-85ss scale bar = 30 µm N = 3 (E-E’’’’) HCR on *tbxt* mRNA (cyan) and *sox2* mRNA (red) Hoechst = nuclear staining (grey) at the 107-120ss scale bar = 30 µm N= 3 (F) average expression of *tbxt* across different somitogenesis stages normalised to percentage of PSM length 16-22ss (green) N= 2, 33-36ss (pink) N= 2, 42-47ss (orange) N = 3, 75-85ss (blue) N= 3, 107-120ss (red) N = 3 (F’) average expression of *sox2* across different somitogenesis stages normalised to percentage of PSM length 16-22ss (green) N = 2, 33-36ss (pink) N = 2, 42-47ss (orange) N = 3, 75-85ss (blue) N = 3, 107-120ss (red) N = 3(G) Brightfield imaging of 44 somite stage control non-amputated embryo at 0hpa hpa = hours post amputation scale bar = 60 µm N= 6 (G’) Brightfield imaging of 75 somite stage control embryo at 16hpa hpa = hours post amputation scale bar = 190 µm N = 3 (H) Brightfield imaging of 44 somite stage whole PSM amputated embryo at 0hpa hpa = hours post amputation scale bar = 60 µm N = 9 (H’) Brightfield imaging of whole PSM amputated embryo with 47 somites at 16hpa hpa = hours post amputation scale bar = 190 µm N = 9 (I) Brightfield imaging of 47 somite stage tailbud posterior PSM amputated embryo at 0hpa hpa = hours post amputation scale bar = 60 µm N= 10 (I’) Brightfield imaging of tailbud posterior PSM amputated embryo with 76 somites at 16hpa hpa = hours post amputation scale bar = 190 µm N = 10 (J) zoomed-in view on control PSM at 16hpa from panel (G’) scale bar = 190 µm (K) zoomed-in view on whole PSM amputation at 16hpa from panel (H’) scale bar = 190 µm (L) zoomed-in view on tailbud posterior PSM amputation at 16hpa from panel (I’) scale bar = 190 µm.

Fig. S8. microsurgical amputation and stemness factors in the PSM

(A) Δ somites = number of somites added at 16hpa in combined data from experiments 1 and 2 for control 30.9 (SD ± 1.16) N = 6 whole PSM amputation 3.8 (SD ± 1.05) N = 15 and posterior PSM amputation 27.1 (SD ± 2.9) N = 14 hpa = hours post amputation SD = standard deviation (B) Brightfield imaging of control non-amputated embryo at 34hpa hpa = hours post amputation scale bar = 200 µm N = 8 (C) Brightfield imaging of tailbud (posterior PSM) amputation embryo at 34hpa hpa = hours post amputation scale bar = 200 µm N = 6 (D) number of somites in control non-amputated embryo at 34hpa 110.38 (SD ± 4.47) and in tailbud (posterior PSM) amputated embryos 107 (SD ± 5.44) hpa = hours post amputation SD = standard deviation Welch two sample t-test p = 0.25 N = 8 control embryos N = 6 tailbud amputated embryos (E-E’’’’) HCR on *pou5f3* mRNA (green) and *gdf11* mRNA (magenta) Hoechst = nuclear staining (grey) at the 16-22ss scale bar = 50 µm N = 4 embryos (F-F’’’’) HCR on *pou5f3* mRNA (green) and *gdf11* mRNA (magenta) Hoechst = nuclear staining (grey) at the 33-36ss scale bar = 30 µm N = 3 embryos (G-G’’’’) HCR on *pou5f3* mRNA (green) and *gdf11* mRNA (magenta) Hoechst = nuclear staining (grey) at the 42-47ss scale bar = 30 µm N = 2 embryos (H-H’’’’) HCR on *pou5f3* mRNA (green) and *gdf11* mRNA (magenta) Hoechst = nuclear staining (grey) at the 75-85ss scale bar = 30 µm N = 3 embryos (I-I’’’’) HCR on *pou5f3* mRNA (green) and *gdf11* mRNA (magenta) Hoechst = nuclear staining (grey) at the 107-120ss scale bar = 30 µm N= 1 embryo (J) average expression of *pou5f3* across different somitogenesis stages normalised to percentage of PSM length 16-22ss (green) N = 4, 33-36ss (pink) N = 3, 42-47ss (orange) N = 2, 75-85ss (blue) N = 3, 107-120ss (red) N = 1 (J’) average expression of *gdf11* across different somitogenesis stages normalised to percentage of PSM length 16-22ss (green) N = 4, 33-36ss (pink) N = 3, 42-47ss (orange) N = 2, 75-85ss (blue) N = 3, 107-120ss (red) N = 1.

Supplementary Fig. S9. HCR on *lin28a/b* and *Hox13* cluster

(A-A’’’’) HCR on *lin28a* mRNA (green) and *lin28b* mRNA (magenta) Hoechst = nuclear staining (grey) at the 16-22ss scale bar = 50 µm N = 2 embryos (B-B’’’’) HCR on *lin28a* mRNA (green) and *lin28b* mRNA (magenta) Hoechst = nuclear staining (grey) at the 33-36ss scale bar = 50 µm N = 3 embryos (C-C’’’’) HCR on *lin28a* mRNA (green) and *lin28b* mRNA (magenta) Hoechst = nuclear staining (grey) at the 42-47ss scale bar = 30 µm N = 2 embryos (D-D’’’’) HCR on *lin28a* mRNA (green) and *lin28b* mRNA (magenta) Hoechst = nuclear staining (grey) at the 75-85ss scale bar = 30 µm N = 2 embryos (E-E’’’’) HCR on *lin28a* mRNA (green) and *lin28b* mRNA (magenta) Hoechst = nuclear staining (grey) at the 107-120ss scale bar = 30 µm N = 5 embryos (F) average expression of *lin28a* across different somitogenesis stages normalised to percentage of PSM length, 16-22ss (green) N = 2, 33-36ss (pink) N = 3, 42-47ss (orange) N = 2, 75-85ss (blue) N = 2, 107-120ss (red) N = 5 (F’) average expression of *lin28b* across different somitogenesis stages normalised to percentage of PSM length 16-22ss (green) N = 2, 33-36ss (pink) N = 3, 42-47ss (orange) N = 2, 75-85ss (blue) N = 2, 107-120ss (red) N = 5 (G-G’’) HCR on *hoxb13a* mRNA (blue) Hoechst = nuclear staining (grey) at the 75-85ss scale bar = 100 µm N = 5 embryos, same embryo shown as in Figure S2C (H-H’’) HCR on *hoxb13a* mRNA (blue) Hoechst = nuclear staining (grey) at the 107-120ss scale bar = 100 µm N= 5 embryos, same embryo shown as in Figure S2D.

## Supplementary Movies legends

### Movie S1

3D reconstructed volumes of 16-22 somite stage N= 9, 33-36 somite stage N= 8, 42-47 somite stage N= 10, 75-85 somite stage N= 8, 107-120 somite stage N= 9 *A. japonica* eels, unsegmented presomitic mesoderm (PSM) = yellow, body axis = purple scale bar = bottom left corner.

### Movie S2

Segmented body axis of a 107-120 somite stage *A. japonica* eel Hoechst = nuclear staining (grey) scale bar = bottom left corner.

### Movie S3

*in-vivo* 4D live confocal imaging of body axis segmentation in somite stage 33-36 *A. japonica* eel embryo injected with H2A-mcherry left panel = bright-field, middle panel = H2A-mcherry (grey), right panel = merged view scale bar = 100 µm, top left= time in hours temperature = 26°C N= 3

### Movie S4

*in-vivo* 4D live confocal imaging of body axis segmentation in somite stage 42-47 *A. japonica* eel embryo injected with H2A-mcherry, left panel = bright-field, middle panel = H2A-mcherry (grey), right panel = merged view scale bar = 100 µm, top left= time in hours temperature = 26°C N= 7.

### Movie S5

3D reconstructed expression of *lfng* mRNA (yellow) in tail (grey) of different somitogenesis stages 16-22 somite stage N= 2, 33-36 somite stage N= 2, 42-47 somite stage N= 3, 75-85 somite stage N= 3, 107-120 somite stage N= 2 *A. japonica* eel scale bar = bottom left.

### Movie S6

3D reconstructed expression of *fgf8a* mRNA (red) in tail (grey) of different somitogenesis stages 16-22 somite stage N= 2, 33-36 somite stage N= 2, 42-47 somite stage N= 2, 75-85 somite stage N= 3, 107-120 somite stage N= 3 *A. japonica* eel scale bar = bottom left.

### Movie S7

3D reconstructed colocalised signal of *sox2 tbxt* mRNA expression (magenta) Hoechst = nuclear staining (grey) at different somitogenesis stages 16-22 somite stage N= 2, 33-36 somite stage N= 2, 42-47 somite stage N= 3, 75-85 somite stage N= 3, 107-120 somite stage N= 3 *A. japonica* eel scale bar = bottom left.

## References and Notes

1. E. N. Rittmeyer, A. Allison, M. C. Gründler, D. K. Thompson, C. C. Austin, Ecological Guild Evolution and the Discovery of the World’s Smallest Vertebrate. PLoS ONE 7, e29797 (2012).

2. J. G. Nielsen, D. G. Smith, The eel family Nemichthyidae (Pisces: Anguilliformes). Dana Report 88, 1–71 (1978).

3. I. Palmeirim, D. Henrique, D. Ish-Horowicz, O. Pourquié, Avian hairy Gene Expression Identifies a Molecular Clock Linked to Vertebrate Segmentation and Somitogenesis. Cell 91, 639–648 (1997).

4. A. Aulehla, W. Wiegraebe, V. Baubet, M. B. Wahl, C. Deng, M. Taketo, M. Lewandoski, O. Pourquié, A β-catenin gradient links the clock and wavefront systems in mouse embryo segmentation. Nat. Cell Biol. 10, 186–193 (2008).

5. C. Gomez, E. M. Özbudak, J. Wunderlich, D. Baumann, J. Lewis, O. Pourquié, Control of segment number in vertebrate embryos. Nature 454, 335–339 (2008).

6. C. Schröter, A. C. Oates, Segment Number and Axial Identity in a Segmentation Clock Period Mutant. Current Biology 20, 1254–1258 (2010).

7. C. Schröter, L. Herrgen, A. Cardona, G. J. Brouhard, B. Feldman, A. C. Oates, Dynamics of zebrafish somitogenesis. Developmental Dynamics 237, 545–553 (2008).

8. P. P. L. Tam, The control of somitogenesis in mouse embryos. Development 65, 103–128 (1981).

9. A. Seleit, I. Brettell, T. Fitzgerald, C. Vibe, F. Loosli, J. Wittbrodt, K. Naruse, E. Birney, A. Aulehla, Modular control of vertebrate axis segmentation in time and space. EMBO J 43, 4068–4091 (2024).

10. H. Suzuki, T. Tanaka, K. Gen, K. Nomura, Y. Kazeto, Optimization of a protocol for fertilized egg production in Japanese eel using recombinant gonadotropins, LHRHa, pimozide, and 17α-hydroxyprogesterone. Aquaculture Reports 37, 102270 (2024).

11. Y. Kazeto, R. Ito, T. Tanaka, H. Suzuki, Y. Ozaki, K. Okuzawa, K. Gen, Establishment of cell-lines stably expressing recombinant Japanese eel follicle-stimulating hormone and luteinizing hormone using CHO-DG44 cells: fully induced ovarian development at different modes. Front. Endocrinol. 14, 1201250 (2023).

12. K. Tsukamoto, Oceanic migration and spawning of anguillid eels. Journal of Fish Biology 74, 1833–1852 (2009).

13. T. Kawakami, Y. Yamada, S. Tanaka, K. Tsukamoto, Prolongation of somitogenesis in two anguilliform species, the Japanese eel *Anguilla japonica* and pike eel *Muraenesox cinereus*, with refined descriptions of their early development. Journal of Fish Biology 90, 1533–1547 (2017).

14. B. Steventon, F. Duarte, R. Lagadec, S. Mazan, J.-F. Nicolas, E. Hirsinger, Species-specific contribution of volumetric growth and tissue convergence to posterior body elongation in vertebrates. Development, 143, 1732–1741 (2016).

15. J. M. Smith, R. Burian, S. Kauffman, P. Alberch, J. Campbell, B. Goodwin, R. Lande, D. Raup, L. Wolpert, Developmental Constraints and Evolution: A Perspective from the Mountain Lake Conference on Development and Evolution. The Quarterly Review of Biology 60, 265–287 (1985).

16. P. Alberch, The logic of monsters: Evidence for internal constraint in development and evolution. Geobios 22, 21–57 (1989).

17. A. B. Ward, R. S. Mehta, Axial Elongation in Fishes: Using Morphological Approaches to Elucidate Developmental Mechanisms in Studying Body Shape. Integrative and Comparative Biology 50, 1106–1119 (2010).

18. R. S. Mehta, A. B. Ward, M. E. Alfaro, P. C. Wainwright, Elongation of the Body in Eels. Integrative and Comparative Biology 50, 1091–1105 (2010).

19. T. Iwamatsu, Stages of normal development in the medaka Oryzias latipes. Mechanisms of Development 121, 605–618 (2004).

20. C. B. Kimmel, W. W. Ballard, S. R. Kimmel, B. Ullmann, T. F. Schilling, Stages of embryonic development of the zebrafish. Developmental Dynamics 203, 253–310 (1995).

21. H. M. T. Choi, M. Schwarzkopf, M. E. Fornace, A. Acharya, G. Artavanis, J. Stegmaier, A. Cunha, N. A. Pierce, Third-generation *in situ* hybridization chain reaction: multiplexed, quantitative, sensitive, versatile, robust. Development 145, dev165753 (2018).

22. M. Gajewski, H. Elmasri, M. Girschick, D. Sieger, C. Winkler, Comparative analysis of her genes during fish somitogenesis suggests a mouse/chick-like mode of oscillation in medaka. Dev Genes Evol 216, 315–332 (2006).

23. M. F. Simsek, A. S. Chandel, D. Saparov, O. Q. H. Zinani, N. Clason, E. M. Özbudak, Periodic inhibition of Erk activity drives sequential somite segmentation. Nature 613, 153–159 (2023).

24. M. F. Simsek, D. Saparov, K. Keseroglu, O. Zinani, A. S. Chandel, B. Dulal, B. K. Sharma, S. Zimik, E. M. Özbudak, The vertebrate segmentation clock drives segmentation by stabilizing Dusp phosphatases in zebrafish. Developmental Cell 60, 669–678.e6 (2025).

25. Y. Niwa, H. Shimojo, A. Isomura, A. González, H. Miyachi, R. Kageyama, Different types of oscillations in Notch and Fgf signaling regulate the spatiotemporal periodicity of somitogenesis. Genes Dev. 25, 1115–1120 (2011).

26. L. Thomson, L. Muresan, B. Steventon, The zebrafish presomitic mesoderm elongates through compaction-extension. Cells & Development 168, 203748 (2021).

27. K. Ishimatsu, T. W. Hiscock, Z. M. Collins, D. W. K. Sari, K. Lischer, D. L. Richmond, Y. Bessho, T. Matsui, S. G. Megason, Size-reduced embryos reveal a gradient scaling based mechanism for zebrafish somite formation. Development, dev.161257 (2018).

28. F. Koch, M. Scholze, L. Wittler, D. Schifferl, S. Sudheer, P. Grote, B. Timmermann, K. Macura, B. G. Herrmann, Antagonistic Activities of Sox2 and Brachyury Control the Fate Choice of Neuro-Mesodermal Progenitors. Developmental Cell 42, 514–526.e7 (2017).

29. S. Edri, P. Hayward, W. Jawaid, A. Martinez Arias, Neuro-mesodermal progenitors (NMPs): a comparative study between pluripotent stem cells and embryo-derived populations. Development 146, dev180190 (2019).

30. A. S. Tucker, J. M. W. Slack, The *Xenopus laevis* tail-forming region. Development 121, 249–262 (1995).

31. R. Bellairs, “The Tail Bud and Cessation of Segmentation in the Chick Embryo” in Somites in Developing Embryos, R. Bellairs, D. A. Ede, J. W. Lash, Eds. (Springer US, Boston, MA, 1986; http://link.springer.com/10.1007/978-1-4899-2013-3_13), pp. 161–178.

32. R. Aires, A. D. Jurberg, F. Leal, A. Nóvoa, M. J. Cohn, M. Mallo, Oct4 Is a Key Regulator of Vertebrate Trunk Length Diversity. Developmental Cell 38, 262–274 (2016).

33. T. Yuikawa, M. Ikeda, S. Tsuda, S. Saito, K. Yamasu, Involvement of Oct4-type transcription factor Pou5f3 in posterior spinal cord formation in zebrafish embryos. Dev Growth Differ 63, 306–322 (2021).

34. A. V. Sánchez-Sánchez, E. Camp, A. García-España, A. Leal-Tassias, J. L. Mullor, Medaka Oct4 is expressed during early embryo development, and in primordial germ cells and adult gonads. Developmental Dynamics 239, 672–679 (2010).

35. R. Aires, L. De Lemos, A. Nóvoa, A. D. Jurberg, B. Mascrez, D. Duboule, M. Mallo, Tail Bud Progenitor Activity Relies on a Network Comprising Gdf11, Lin28, and Hox13 Genes. Developmental Cell 48, 383–395.e8 (2019).

36. D. A. Robinton, J. Chal, E. Lummertz Da Rocha, A. Han, A. V. Yermalovich, M. Oginuma, T. M. Schlaeger, P. Sousa, A. Rodriguez, A. Urbach, O. Pourquié, G. Q. Daley, The Lin28/let-7 Pathway Regulates the Mammalian Caudal Body Axis Elongation Program. Developmental Cell 48, 396–405.e3 (2019).

37. H. Miyazawa, Y. Muramatsu, H. Makino, Y. Yamaguchi, M. Miura, Temporal regulation of *Lin28a* during mammalian neurulation contributes to neonatal body size control. Developmental Dynamics 248, 931–941 (2019).

38. Z. Ye, D. Kimelman, Hox13 genes are required for mesoderm formation and axis elongation during early zebrafish development. Development 147, dev185298 (2020).

39. T. J. Near, R. I. Eytan, A. Dornburg, K. L. Kuhn, J. A. Moore, M. P. Davis, P. C. Wainwright, M. Friedman, W. L. Smith, Resolution of ray-finned fish phylogeny and timing of diversification. Proc. Natl. Acad. Sci. U.S.A. 109, 13698–13703 (2012).

40. P. J. Bergmann, G. Morinaga, The convergent evolution of snake-like forms by divergent evolutionary pathways in squamate reptiles*. Evolution 73, 481–496 (2019).

41. P. Alberch, S. J. Gould, G. F. Oster, D. B. Wake, Size and Shape in Ontogeny and Phylogeny. Paleobiology 5, 296–317 (1979).

42. S. J. Gould, Ontogeny and Phylogeny (The Belknap Press of Harvard Univ. Press, Cambridge, Mass., 16. print., 2002).

43. H. Ohta, Y. Sato, H. Imaizumi, Y. Kazeto, Changes in milt volume and sperm quality with time after an injection of recombinant Japanese eel luteinizing hormone in male Japanese eels. Aquaculture 479, 150–154 (2017).

44. K. Nomura, I. C. C. Koh, R. Iio, D. Okuda, Y. Kazeto, H. Tanaka, H. Ohta, Sperm cryopreservation protocols for the large-scale fertilization of Japanese eel using a combination of large-volume straws and low sperm dilution ratio. Aquaculture 496, 203–210 (2018).

45. H. Ohta, H. Kagawa, H. Tanaka, T. Unuma, Control by the environmental concentration of ions of the potential for motility in Japanese eel spermatozoa. Aquaculture 198, 339–351 (2001).

46. H. S Bruce, G. Jerz, S. R Kelly, J. McCarthy, A. Pomerantz, G. Senevirathne, A. Sherrard, D. A Sun, C. Wolff, N. H Patel, Hybridization Chain Reaction (HCR) In Situ Protocol v1. [Preprint] (2021). 10.17504/protocols.io.bunznvf6.

47. E. Kuehn, D. S. Clausen, R. W. Null, B. M. Metzger, A. D. Willis, B. D. Özpolat, Segment number threshold determines juvenile onset of germline cluster expansion in *Platynereis dumerilii*. J Exp Zool Pt B 338, 225–240 (2022).

48. B. M. Kirilenko, C. Munegowda, E. Osipova, D. Jebb, V. Sharma, M. Blumer, A. E. Morales, A.-W. Ahmed, D.-G. Kontopoulos, L. Hilgers, K. Lindblad-Toh, E. K. Karlsson, Zoonomia Consortium‡, M. Hiller, G. Andrews, J. C. Armstrong, M. Bianchi, B. W. Birren, K. R. Bredemeyer, A. M. Breit, M. J. Christmas, H. Clawson, J. Damas, F. Di Palma, M. Diekhans, M. X. Dong, E. Eizirik, K. Fan, C. Fanter, N. M. Foley, K. Forsberg-Nilsson, C. J. Garcia, J. Gatesy, S. Gazal, D. P. Genereux, L. Goodman, J. Grimshaw, M. K. Halsey, A. J. Harris, G. Hickey, M. Hiller, et al., Integrating gene annotation with orthology inference at scale. Science 380, eabn3107 (2023).

49. N. Hecker, M. Hiller, A genome alignment of 120 mammals highlights ultraconserved element variability and placenta-associated enhancers. GigaScience 9, giz159 (2020).

50. V. Sharma, M. Hiller, Increased alignment sensitivity improves the usage of genome alignments for comparative gene annotation. Nucleic Acids Research 45, 8369–8377 (2017).

51. W. J. Kent, R. Baertsch, A. Hinrichs, W. Miller, D. Haussler, Evolution’s cauldron: Duplication, deletion, and rearrangement in the mouse and human genomes. Proc. Natl. Acad. Sci. U.S.A. 100, 11484–11489 (2003).

52. E. Osipova, N. Hecker, M. Hiller, RepeatFiller newly identifies megabases of aligning repetitive sequences and improves annotations of conserved non-exonic elements. GigaScience 8, giz132 (2019).

53. H. G. Suarez, B. E. Langer, P. Ladde, M. Hiller, chainCleaner improves genome alignment specificity and sensitivity. Bioinformatics 33, 1596–1603 (2017).

54. E. Hirsinger, B. Steventon, A Versatile Mounting Method for Long Term Imaging of Zebrafish Development. JoVE, 55210 (2017).

55. J. Schindelin, I. Arganda-Carreras, E. Frise, V. Kaynig, M. Longair, T. Pietzsch, S. Preibisch, C. Rueden, S. Saalfeld, B. Schmid, J.-Y. Tinevez, D. J. White, V. Hartenstein, K. Eliceiri, P. Tomancak, A. Cardona, Fiji: an open-source platform for biological-image analysis. Nat Methods 9, 676–682 (2012).

56. J. Goedhart, PlotTwist: A web app for plotting and annotating continuous data. PLoS Biol 18, e3000581 (2020).

57. M. Postma, J. Goedhart, PlotsOfData—A web app for visualizing data together with their summaries. PLoS Biol 17, e3000202 (2019).

58. C. B. Vibe, The temperature response of the medaka segmentation clock and its link to robustness in embryonic patterning. doi: 10.11588/HEIDOK.00028769 (2020).

