## Supplementary Figures for "Heterochrony of axis segmentation underlies extreme morphogenesis in the Japanese eel"

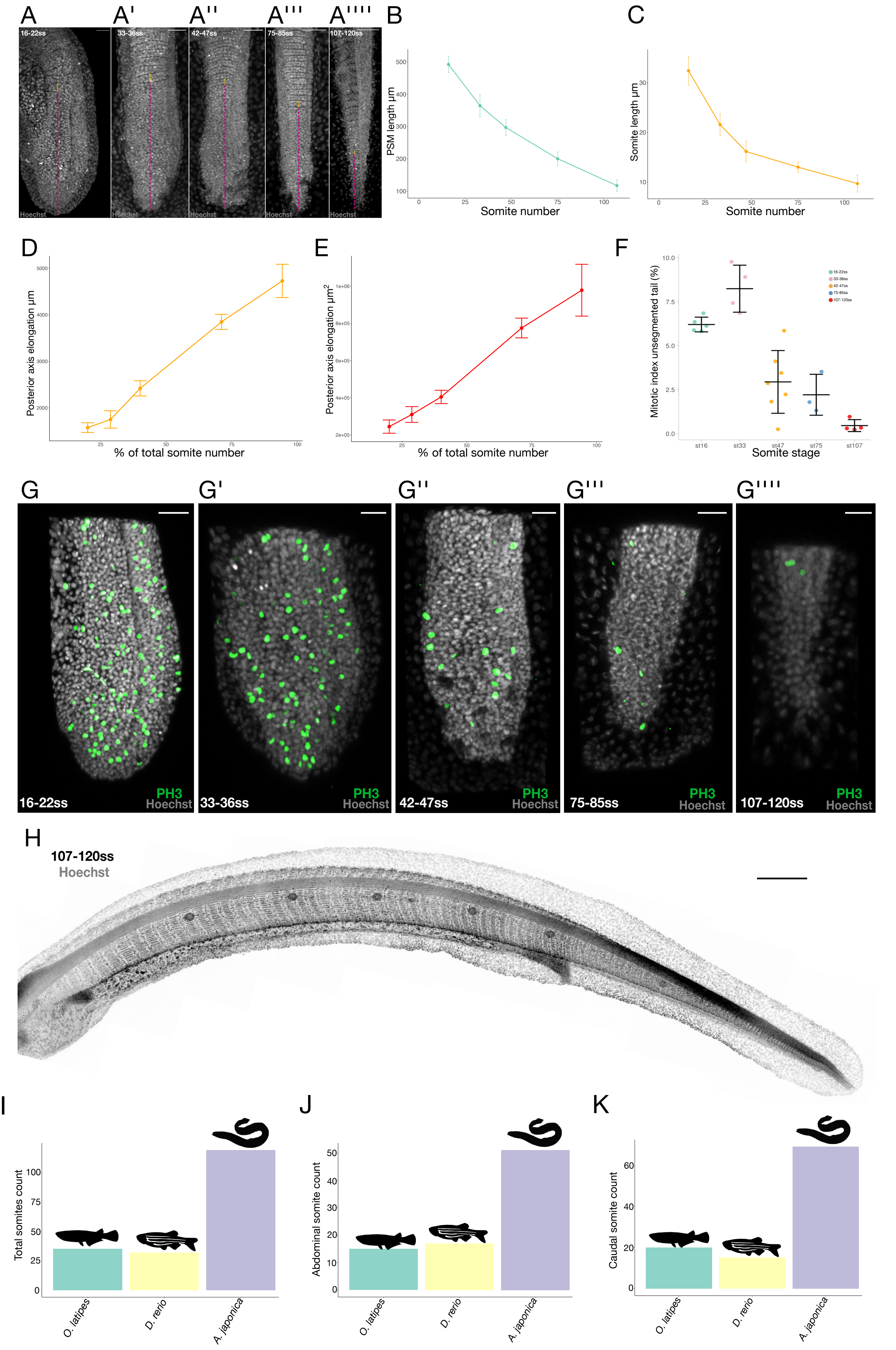


**Seleit et al., Supplementary Figure 1**

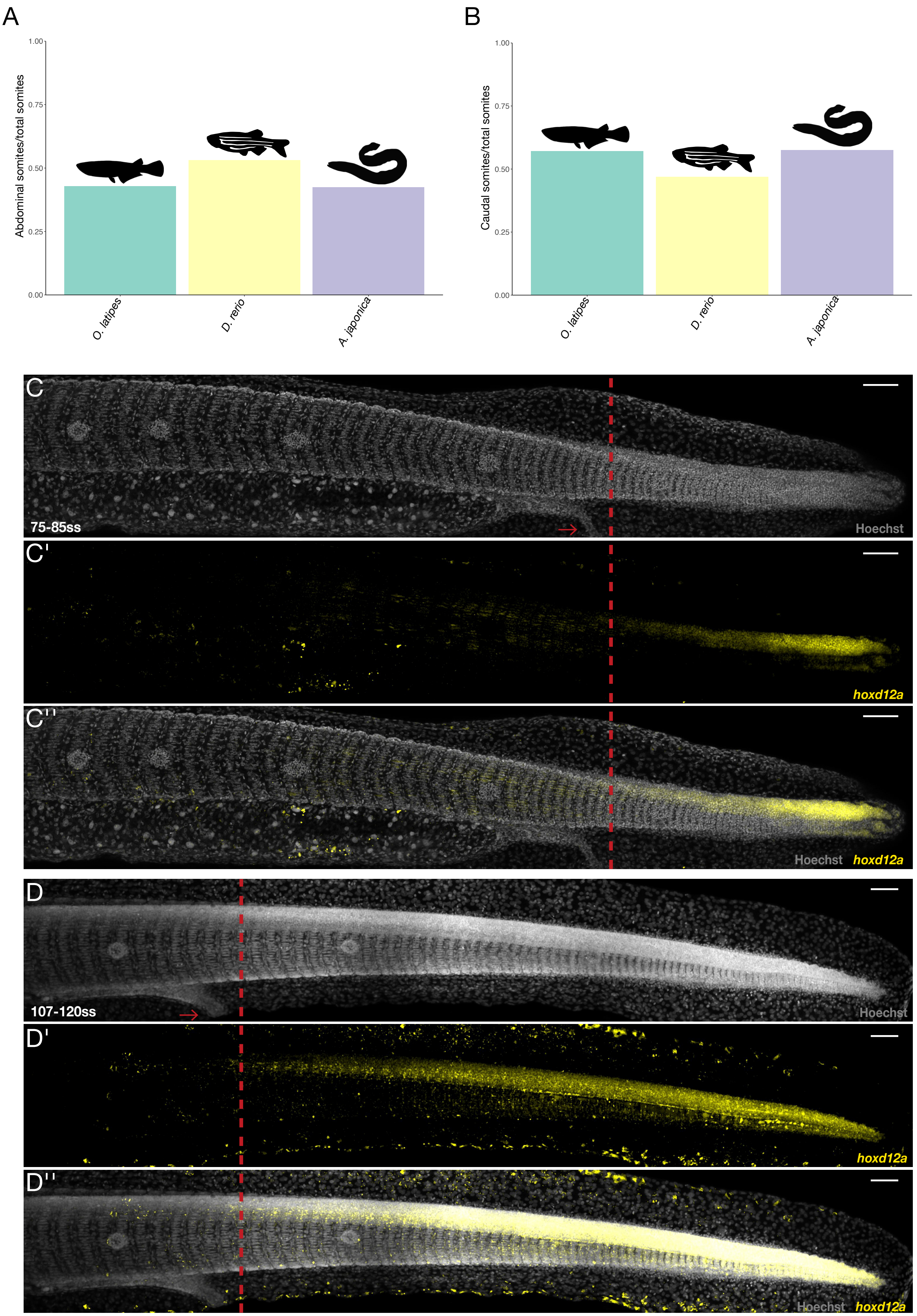


**Seleit et al., Supplementary Figure 2**

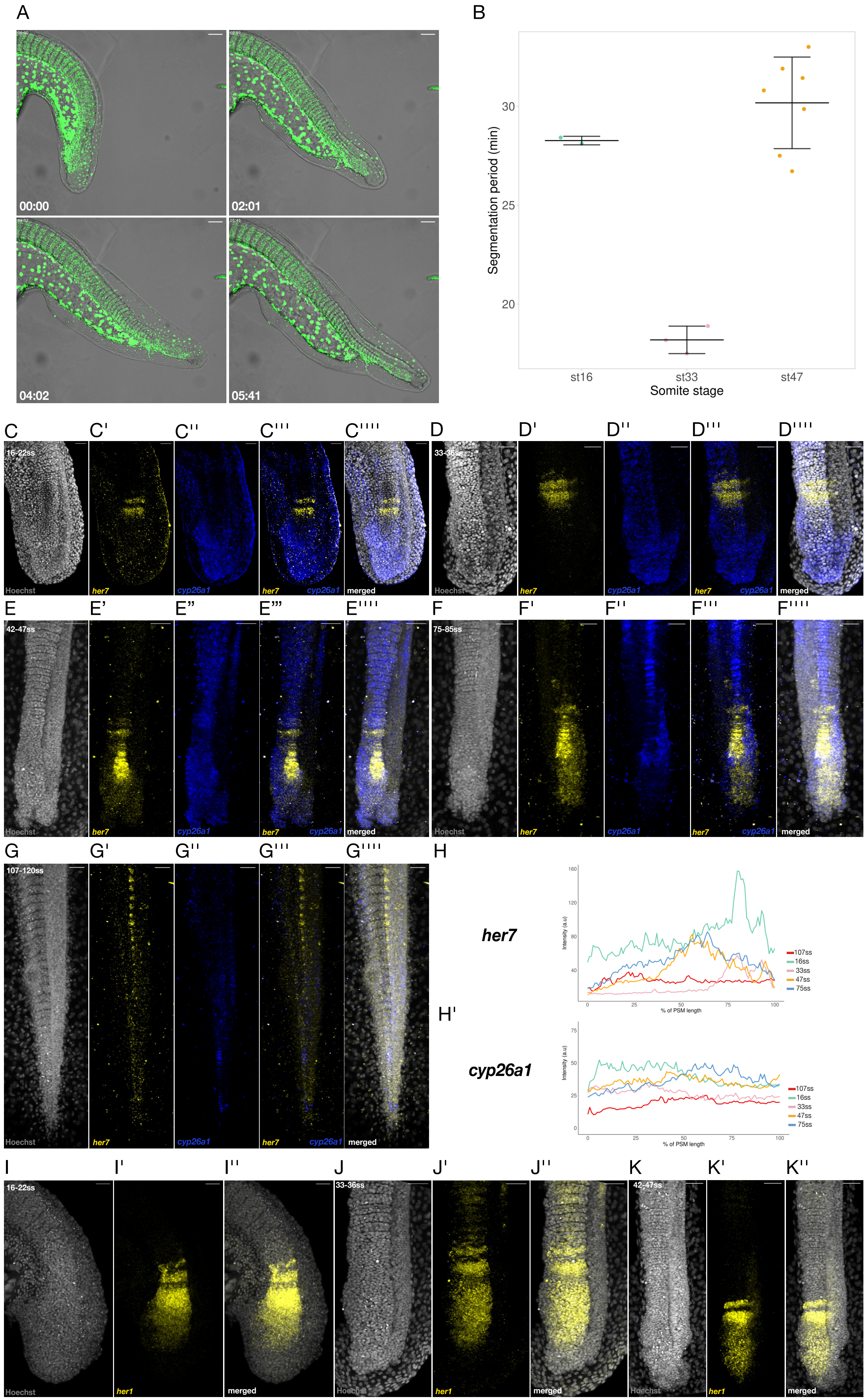


**Seleit et al., Supplementary Figure 3**

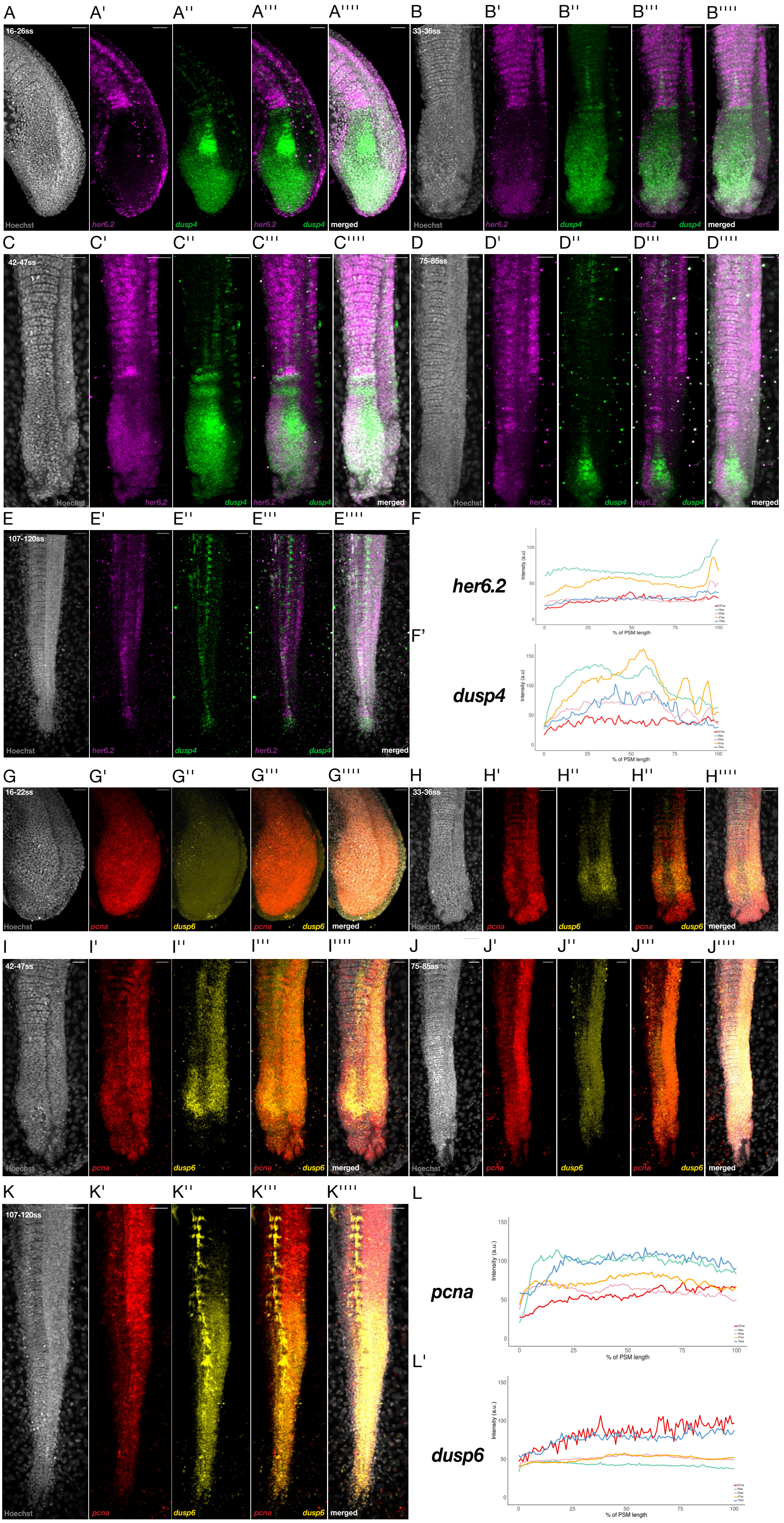


**Seleit et al., Supplementary Figure 4**

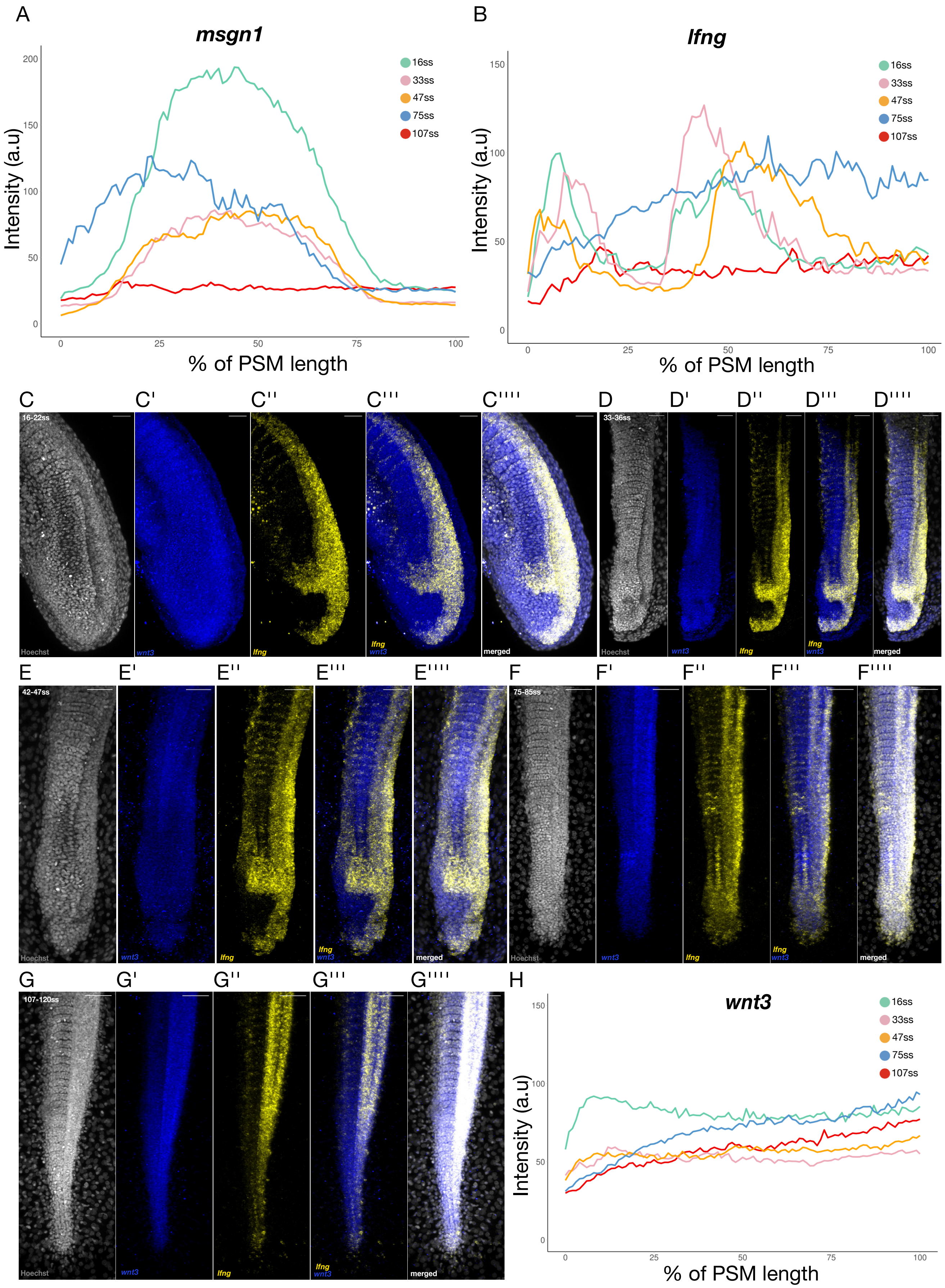


**Seleit et al., Supplementary Figure 5**

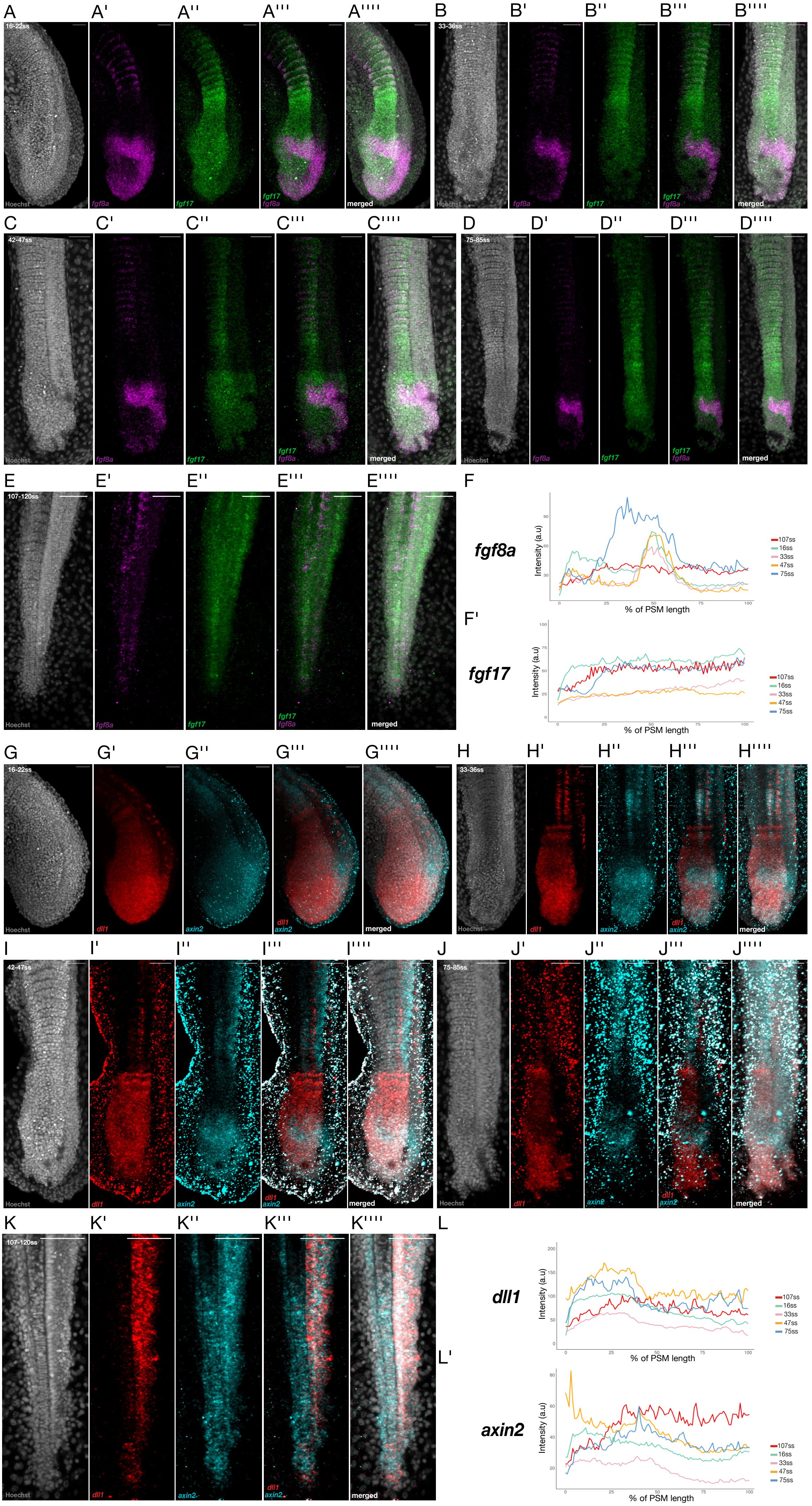


**Seleit et al., Supplementary Figure 6**

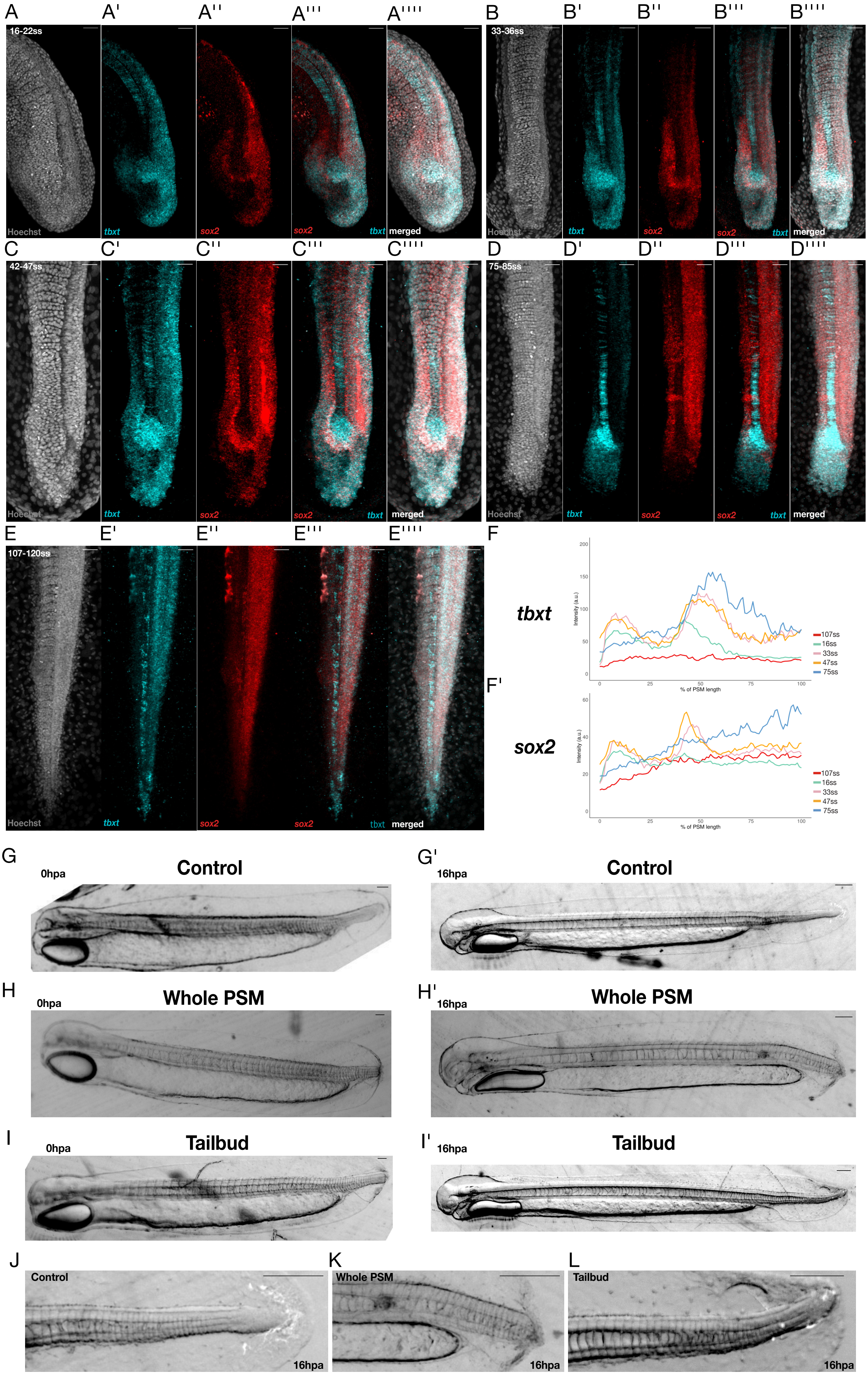


**Seleit et al., Supplementary Figure 7**

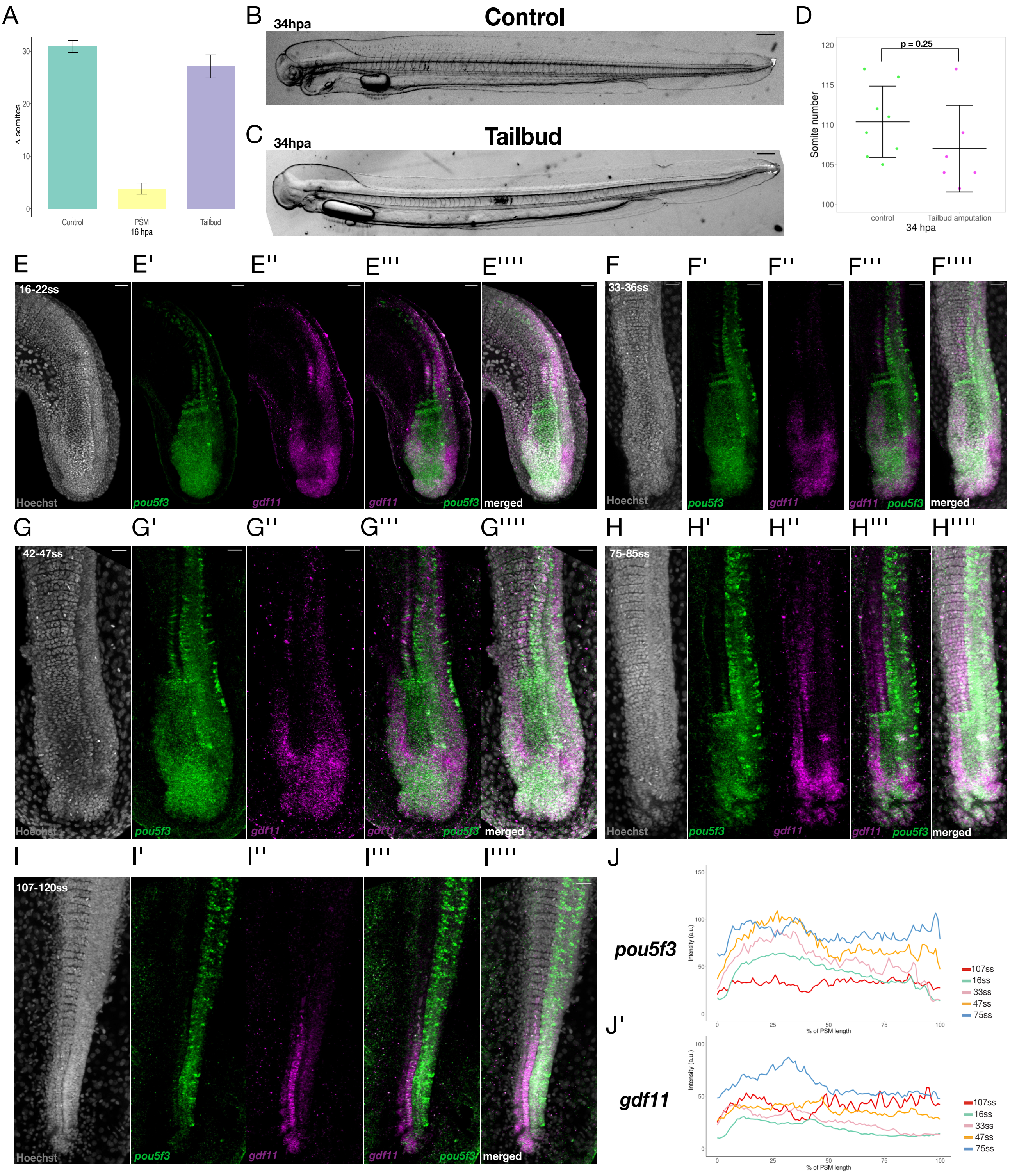


**Seleit et al., Supplementary Figure 8**

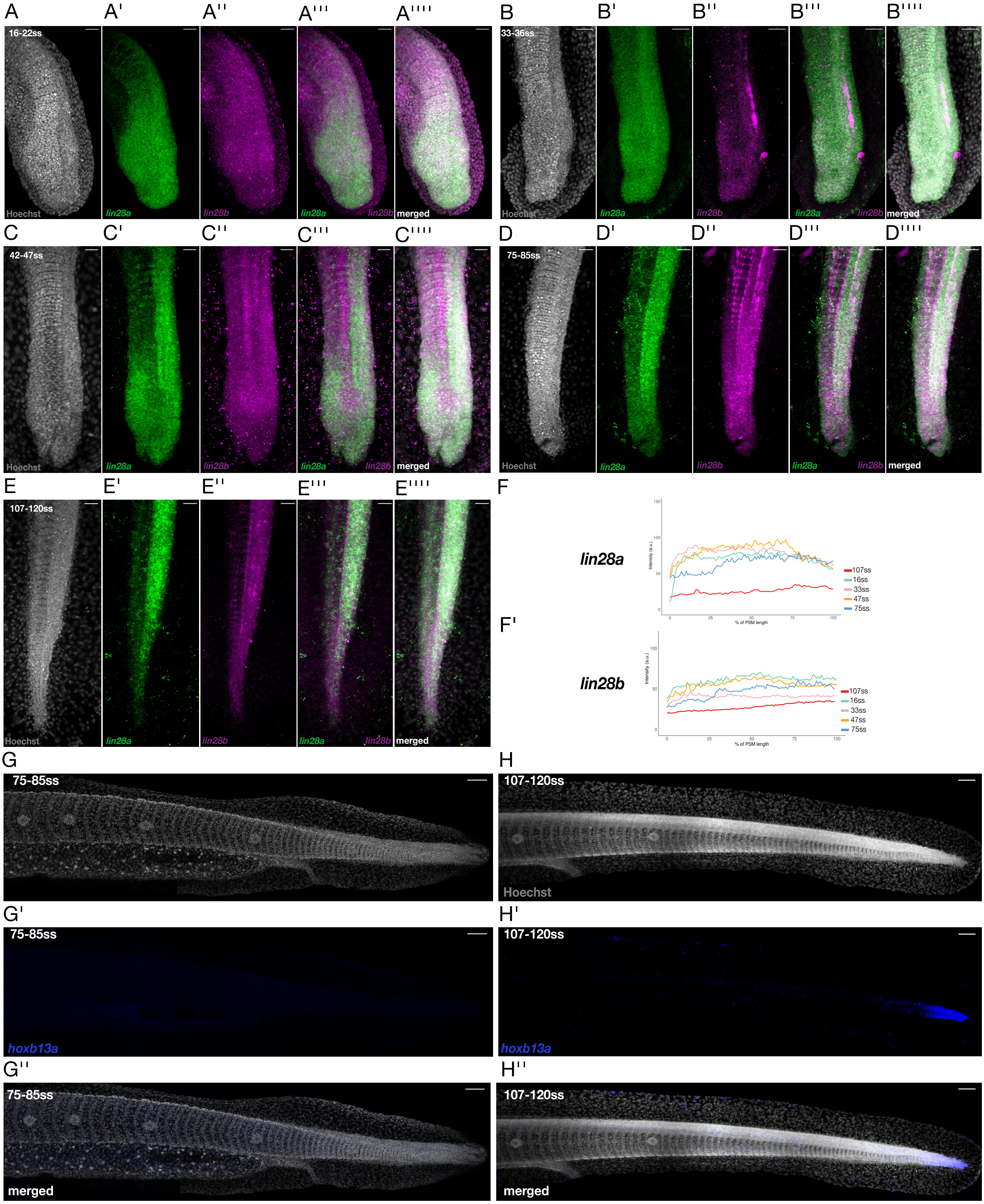


**Seleit et al., Supplementary Figure 9**
